# Structural asymmetry governs the assembly and GTPase activity of McrBC restriction complexes

**DOI:** 10.1101/2020.01.30.927467

**Authors:** Yiming Niu, Hiroshi Suzuki, Christopher J. Hosford, Thomas Walz, Joshua S. Chappie

**Author notes:** To whom correspondence should be addressed (T.W.), (J.S.C.). These authors contributed equally and are listed in alphabetical order (Yiming Niu, Hiroshi Suzuki).

## Abstract

McrBC complexes are motor-driven nucleases functioning in bacterial self-defense by cleaving foreign DNA. The GTP-specific AAA+ protein McrB powers translocation along DNA and its hydrolysis activity is stimulated by its partner nuclease McrC. Here, we report cryo-EM structures of *Thermococcus gammatolerans* McrB and McrBC, and *E. coli* McrBC. The McrB hexamers, containing the necessary catalytic machinery for basal GTP hydrolysis, are intrinsically asymmetric. This asymmetry directs McrC binding so that it engages a single active site, where it then uses an arginine/lysine-mediated hydrogen-bonding network to reposition the asparagine in the McrB signature motif for optimal catalytic function. While the two McrBC complexes use different DNA-binding domains, these contribute to the same general GTP-recognition mechanism employed by all G proteins. Asymmetry also induces distinct inter-subunit interactions around the ring, suggesting a coordinated and directional GTP-hydrolysis cycle. Our data provide novel insights into the conserved molecular mechanisms governing McrB family AAA+ motors.

## INTRODUCTION

Infections by antibiotic-resistant bacteria pose a serious threat to human health (Resistance, 2016; Solomon and Oliver, 2014). The slow progress in developing new drugs to combat these emerging ‘superbugs’ and the rapid exchange of resistance genes among microbial populations has intensified the need for alternative therapeutic strategies (Theuretzbacher and Piddock, 2019). One such strategy employs bacteriophages (phages) – viruses that infect a bacterial host, replicate, and then induce cell lysis to release the mature phage progeny, killing the host in the process (Kortright et al., 2019). The pharmaceutical application of phages dates back to the early 1920s (Wittebole et al., 2014) and has resurged in recent years, bolstered by success in a number of clinical settings (Dedrick et al., 2019; Schooley et al., 2017). Despite these promising results, phage therapy faces numerous challenges. One significant hurdle is that bacteria have evolved an array of defense mechanisms, including restriction modification systems, modification-dependent restrictions systems (MDRS), phage-exclusion systems, and CRISPR-Cas adaptive immune systems, that can hinder phage infection and diminish their subsequent killing potential (Hille et al., 2018; Labrie et al., 2010). These machineries lack eukaryotic homologs and are conserved across antibiotic-resistant bacteria like methicillin-resistant *Staphylococcus aureus* (MRSA), *Clostridium difficile*, and *Klebsiella pneumoniae*, making their components promising candidates for targeted inhibition. Some phages indeed already encode inhibitor proteins that can neutralize restriction and/or CRISPR systems (Samson et al., 2013; Stanley and Maxwell, 2018), allowing them to survive and kill under conditions in which they would normally be suppressed. Elucidating the structure and function of bacterial defense systems will therefore extend these principles and aid in the development of new drugs that increase phage efficacy.

McrBC is a two-component MDRS that in *E. coli* (Ec) restricts phage DNA and foreign DNA containing methylated cytosines (Luria and Human, 1952; Weigele and Raleigh, 2016). EcMcrB consists of an N-terminal DNA-binding domain that targets fully or hemi-methylated R^M^C sites (where R is a purine base and ^M^C is a 4-methyl-, 5-methyl- or 5-hydroxymethyl-cytosine) (Gast et al., 1997; Kruger et al., 1995; Pieper et al., 1999b; Sukackaite et al., 2012; Sutherland et al., 1992; Zagorskaite et al., 2018) and a C-terminal AAA+ (extended <u>A</u>TPases Associated with various cellular Activities) domain that hydrolyzes GTP and oligomerizes into hexamers (Nirwan et al., 2019a; Panne et al., 2001). EcMcrB’s basal GTPase activity (~0.5-1 min^−1^) is stimulated ~30-40-fold *in vitro* via interaction with its partner EcMcrC (Pieper et al., 1999b), a PD-(D/E)xK family endonuclease that cannot stably bind DNA on its own and thus associates with the hexameric McrB AAA+ ring (Panne et al., 2001). Biochemical data suggest that stimulated GTP hydrolysis powers DNA translocation (Panne et al., 1999; Sutherland et al., 1992), allowing EcMcrBC complexes bound to distant R^M^C sites to interact and induce cleavage on both strands (Pieper et al., 2002; Stewart et al., 2000). While these activities have yet to be demonstrated *in vitro* for homologs beyond *E. coli,* other family members have also been shown to function in bacterial defense *in vivo* (O’Driscoll et al., 2006; O’Sullivan et al., 1995; Ohshima et al., 2002). These machines, however, exhibit different specificities for DNA modifications and/or sequences (Hosford and Chappie, 2018; O’Driscoll et al., 2006; Ohshima et al., 2002; Yang et al., 2016, Hosford et al., 2020), suggesting that the core machinery for GTP hydrolysis and DNA cleavage is conserved and has been adapted to different targets throughout evolution in response to various selective pressure from invading phages. This flexibility holds a tremendous potential for engineering new endonucleases for biotechnology and biomedical applications, providing further motivation to study the structural organization and functional regulation of McrBC complexes.

AAA+ proteins are large, multimeric machines that use the energy of ATP hydrolysis to power a wide array of cellular processes (Snider et al., 2008). These enzymes are built around a common structural core (Neuwald et al., 1999) and contain numerous conserved sequence elements important for nucleotide binding and hydrolysis (Erzberger and Berger, 2006). AAA+ protein active sites are formed at the interface between two monomers, thus requiring higher-order assembly – predominantly as hexamers – for function (Wendler et al., 2012). As a consequence, some catalytic residues like charge-compensating arginine fingers are provided in *trans* by the neighboring subunit. Despite sharing a common architecture, McrB is the only AAA+ protein that preferentially binds and hydrolyzes GTP (Pieper et al., 1997; Pieper et al., 1999a). All McrB homologs contain a conserved consensus sequence of MNxxDRS that replaces the AAA+ sensor I motif and is predicted to function as a G4 element, which confers guanine-nucleotide specificity in GTPases (Bourne et al., 1991). Mutation of this segment, however, does not significantly alter the nucleotide-binding profile of *E. coli* McrB (Pieper et al., 1999a), indicating that other regions of the protein dictate GTP selectivity. Stimulation of hydrolysis by a binding partner is also rare among AAA+ proteins but reminiscent of the activation of small GTPases by their corresponding GTPase-activating proteins (GAPs) (Paduch et al., 2001). A key difference, however, is that McrC only exerts its effects on the assembled McrB oligomer. Elucidating the structural basis for GTP recognition and stimulated hydrolysis is important for defining McrBC’s divergence from other members of both the AAA+ and GTPase superfamilies.

A recent cryo-EM reconstruction of the hexameric EcMcrB AAA+ domain in complex with EcMcrC (Nirwan et al., 2019b) provided the first structural view of McrBC but fell short of answering many important mechanistic questions. Here, we present cryo-EM structures of an McrB hexamer and McrBC complexes from the evolutionarily distant archaeal species *Thermococcus gammatolerans* (Tg) and the well-characterized *E. coli* system. Our models confirm that McrBC complexes share the same general architecture but lead to a different view of the GTP hydrolysis cycle wherein structural asymmetry drives the underlying physical interactions and conformational motions. Moreover, our structures provide a detailed molecular mechanism for how McrC binding stimulates McrB GTP hydrolysis, which we show is conserved across the McrBC family. Our structures also establish that McrB homologs use the same general chemistry employed by all GTPases to recognize GTP, albeit through different structural elements upstream of the AAA+ domain. This observation establishes how distant McrB homologs have adapted and maintained guanine-nucleotide specificity despite the individual constraints imposed by their structurally unrelated N-terminal domains. Together these data provide novel insights into the structure, function, and regulation of motor-driven McrBC nucleases.

## RESULTS

### TgMcrB^AAA^ forms an asymmetric hexamer

Given the widespread distribution of *mcrBC* genes among diverse bacteria and archaea, we sought to examine the structural and biochemical properties of different McrB homologs to understand how these AAA+ enzymes have evolved to preferentially bind and hydrolyze GTP. Our previous work identified the archaeal McrB homolog from *Thermococcus gammatolerans* (TgMcrB) as an ideal candidate for structural studies given its compact size and increased thermostability (Hosford et al., 2020). The purified AAA+ domain from TgMcrB (TgMcrB^AAA^) forms stable oligomers even in the absence of nucleotides (Figure S1a). Single-particle cryo-EM analysis of purified TgMcrB^AAA^ incubated with the non-hydrolyzable GTP analog GTPγS yielded a density map at an overall resolution of 3.1 Å with no symmetry imposed (Figures 1 and S1). The cryo-EM map reveals that TgMcrB^AAA^ forms a ring-shaped, homohexameric assembly with six nucleotides bound at the subunit interfaces, similar to the closed-ring assembly seen in Type I AAA ATPases (Gai et al., 2004; Monroe et al., 2017). Each subunit displays a canonical AAA+ fold with the additional features of a β-hairpin inserted in helix 2 of the large subdomain as previously predicted (Iyer et al., 2004) and ‘wing’-like helices in the small subdomain (Supplementary Data S1 and S2). The TgMcrB^AAA^ hexamer is asymmetric with four tight interfaces (between monomers B/C, C/D, D/E, and E/F) that bury a surface area ranging from 2393 to 2554 Å^2^ and two loose interfaces (between monomers A/B and F/A) that bury surface areas of 1519 and 1772 Å^2^ (Figure 1c). Tight interfaces feature a hydrogen bond between Asp420 in one monomer and Arg360 in the adjacent monomer (Figure 1d). Arg414 from the first monomer also extends into the neighboring monomer, where it forms hydrogen bonds with Glu527 and π-stacking interactions with Tyr530 (Figure 1d). These interactions are absent at the loose interfaces, where Glu527 instead interacts *in trans* with Arg424 (Figure 1e). All of these residues are highly conserved amongst McrB family proteins (Supplementary Data S1).

**Figure 1.**
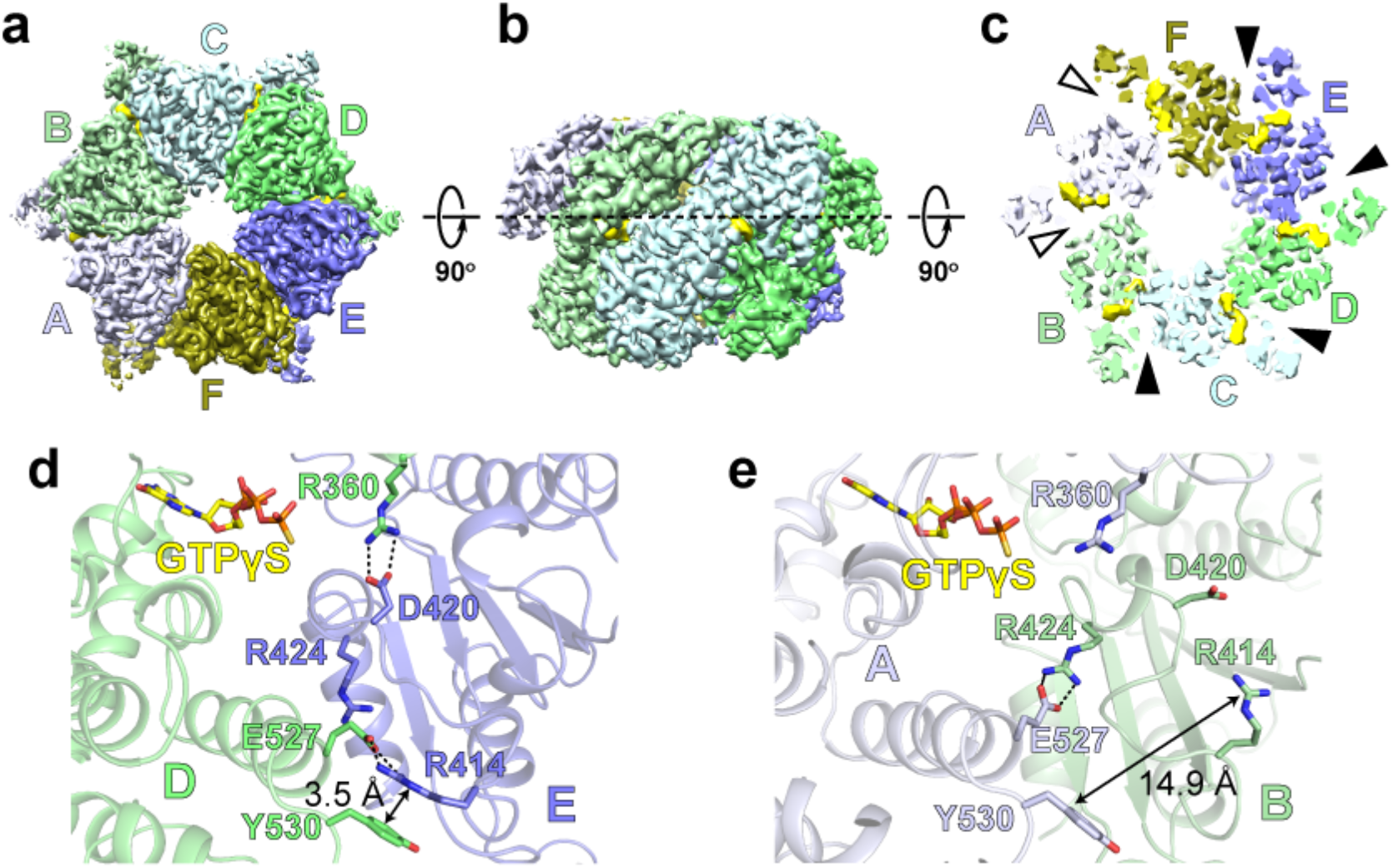
TgMcrB^AAA^ forms an asymmetric hexamer. (**a** and **b**) Bottom and side views of the cryo-EM density map of the TgMcrB^AAA^ hexamer. Subunits are colored in shades of blue and green and nucleotides are shown in yellow. (**c**) Slice section through the TgMcrB^AAA^ hexamer map at the level of the bound nucleotides, indicated by the dashed line in (**b**). Solid and empty arrowheads indicate tight and loose interfaces respectively. (**d** and **e**) Close-up views of interacting residues at the tight D/E interface (**d**) and the loose A/B interface (**e**). Dashed lines indicate hydrogen bonds.

To determine if these interface residues affect McrB’s catalytic turnover, we mutated each side chain individually to alanine in the context of TgMcrB^AAA^ and measured basal GTPase activity using a colorimetric assay. All mutants show an approximate two-fold increase in hydrolysis activity compared to that of the wild-type protein (Figure S1i). Alanine substitution of Arg337 in EcMcrB (corresponding to Arg414 in TgMcrB, Supplementary Data S2) was previously shown to increase the basal GTPase rate three-fold (Pieper et al., 1999a), consistent with our results.

In parallel, we also determined the structure of TgMcrB^AAA^ in the presence of GTPγS by X-ray crystallography at 2.95-Å resolution (Supplementary Table S1). Symmetry-related hexamers abut against each other in the crystal lattice, deforming the planar arrangement of the six subunits in each molecule. This produces an ‘open-ring’ conformation with the subunits at one interface significantly splayed apart and the small subdomain of the A subunit highly disordered (Figure S1j). The individual TgMcrB^AAA^ monomers, however, adopt the same overall conformation and organization of nucleotide binding as is observed in the cryo-EM reconstruction (Figure S1k), suggesting the distorted appearance of the hexamer is an artifact of crystal-packing forces that strain the loose interfaces.

### TgMcrB^AAA^ contains the complete machinery for nucleotide hydrolysis

Nucleotide hydrolases harness the energy of ATP or GTP hydrolysis to catalyze energetically unfavorable biological reactions, coordinate signal-transduction events, and power protein conformational changes that orchestrate a multitude of cellular processes (Vetter and Wittinghofer, 1999). Efficient hydrolysis requires (i) the binding and recognition of the appropriate nucleotide substrate, (ii) the correct positioning of a water molecule for an in-line SN2 attack on the γ-phosphate to initiate cleavage of the phosphoanhydride bond, and (iii) neutralization of a negative charge that develops between the β- and γ-phosphates in the transition state (Chappie and Dyda, 2013).

While a conserved sequence motif of GxxGxGK[T/S] (P-loop/Walker A motif) coordinates the α- and β-phosphates in both ATPases and GTPases (Wittinghofer, 2016), the remaining catalytic machinery, specificity determinants, and charge-compensating elements vary from enzyme to enzyme. AAA+ proteins contain four additional sequence motifs – Walker B, Sensor I, Sensor II, and second region of homology (SRH) – that contribute to ATP binding and hydrolysis along with the conserved P-loop/Walker A motif (Erzberger and Berger, 2006; Miller and Enemark, 2016). The Walker B motif (D[D/E]xx) stabilizes an essential magnesium cofactor and acts in concert with a polar residue in the Sensor I motif to orient the catalytic water for nucleophilic attack on the γ-phosphate. The Sensor II motif localizes to helix 7 and contains a conserved arginine that interacts with the γ-phosphate. By convention, the subunit contributing these structural motifs to the nucleotide-binding pocket is referred to as the *cis* subunit. The neighboring, *trans* subunit inserts the arginine finger at the end of helix 4 of the SRH into the nucleotide-binding pocket, where it stabilizes the γ-phosphate and contributes to the charge compensation in the transition state.

Each composite active site of TgMcrB^AAA^ contains one GTPγS molecule and a bound magnesium ion (Figure 2a-c). In the *cis* subunit, the main-chain atoms of the Walker A motif interact with the α- and β-phosphates and Lys221 contacts the γ-phosphate of GTPγS (Figure 2a-b). Thr222 (Walker A) and Asp356 (Walker B) coordinate the magnesium cofactor along with two ordered water molecules (Figure 2a). Mutation of these conserved side chains to alanine impairs the basal GTPase activity of TgMcrB^AAA^ (Figure 2c-d). Glu537 (Walker B) lies in close proximity to the γ-phosphate, primed to help stabilize a catalytic water (Figure 2a). An alanine substitution at this position completely abolishes hydrolysis activity (Figure 2d). Negative-stain EM indicates that the Asp356Ala mutation has a higher propensity to disrupt the TgMcrB^AAA^ hexamer than the Glu357Ala mutation (Figure 2e), consistent with their distinct functions in nucleotide binding/stabilization versus catalysis. This result mirrors the different effects on oligomerization observed when the corresponding residues (Asp279 and Glu280) were mutated in EcMcrB (Nirwan et al., 2019a). Notably, the conserved McrB consensus loop (^409^MNxxDR^414^) replaces Sensor I and is located close to the γ-phosphate (Figure 2 and Supplementary Data S1). Asn410Ala and Asp413Ala mutants significantly impair basal GTPase activity (Figure 2d), suggesting they are critical for catalytic turnover, rather than for nucleotide binding as was previously predicted (Pieper et al., 1997).

**Figure 2.**
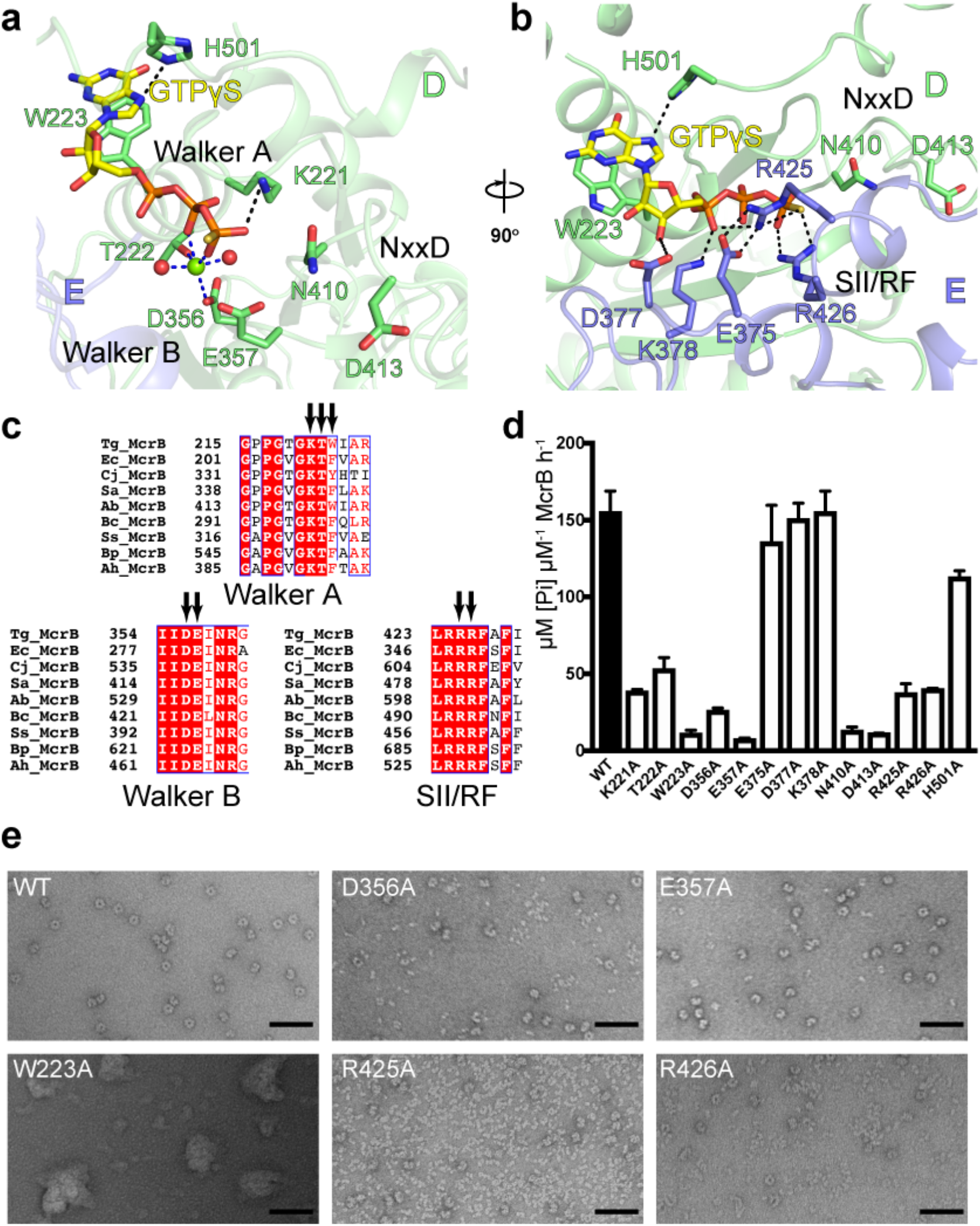
Catalytic residues involved in the basal GTPase activity of TgMcrB. (**a** and **b**) Close-up views of the GTP-binding site at the tight D/E interface, highlighting residues involved in *cis* interactions, in particular those of the Walker A and B motifs and the NxxD motif (**a**), and residues involved in *trans* interactions, in particular those of the Sensor II/arginine finger (SII/RF) motif (**b**). Spheres indicate waters (red) and a magnesium ion (green). Dashed lines indicate hydrogen bonds (black) and metal coordination (blue). (**c**) Sequence alignment of McrB homologs for the classic Walker A and B motifs and the SII/RF motif. Arrows indicate the catalytic residues. Sequence alignment abbreviations are as follows: Tg, *Thermococcus gammatolerans*; Ec, *Escherichia coli*; Cj, *Campylobacter jejuni*; Sa, *Staphylococcus aureus*; Ab, *Aciduliprofundum boonei*; Bc, *Bacillus cereus*; Ss, *Streptococcus suis*; Bp, *Butyrivibrio proteoclasticus*; Ah, *Anaerobutyricum hallii*. (**d**) Basal GTPase activity of wild-type TgMcrB and alanine mutants at the residues shown in (**a**) and (**b**) (n = 3, mean ± standard deviation). (**e**) Selected micrograph areas of negatively stained wild-type and mutant TgMcrB^AAA^. Scale bars are 50 nm.

His501 and Trp223 in the *cis* subunit sandwich the guanine base of GTPγS (Figure 2a). His501 is situated above and forms a hydrogen bond with the 7’ nitrogen. Trp223, which lies adjacent to the Walker A motif, forms a unique parallel π-stacking interaction from below that has never been observed nor predicted for any GTPase or AAA+ protein (Iyer et al., 2004; Leipe et al., 2002). Mutation of Trp223 to Ala completely abolishes the basal GTPase activity (Figure 2d) and causes the protein to aggregate, as seen by negative-stain EM imaging (Figure 2e). These observations indicate that π-stacking is critical for both McrB GTP binding and the stability of the oligomeric assembly. Interestingly, the analogous residue (Phe209) was never mutated in previous studies of EcMcrB as it is not strictly conserved across the McrB family. We do note, however, that every homolog contains a residue at this position that is capable of π-stacking (Trp, Phe, Tyr or Arg) (Figure 2c and Supplementary Data S1).

The *trans* subunit also contributes numerous conserved side chains that stabilize different portions of the bound nucleotide. Asp377 interacts with the 3’ ribose hydroxyl group while Glu375 and Lys378 coordinate the α-phosphate (Figure 2b and Supplementary Data S1). Mutations of these side chains had negligible effects on basal GTPase activity (Figure 2d). Arg426 in helix α11 acts as the charge-compensating arginine finger, here forming hydrogen bonds with the γ-phosphate in the ground state (Figure 2b). A second neighboring arginine located on the same helix, Arg425, assumes the role of the missing Sensor II motif (Figure 2b and c). Arg425Ala and Arg426Ala mutations impair basal GTPase activity and disrupt hexamer formation (Figure 2d and e). All the *trans* interactions with GTPγS are prominent at the tight interfaces but are lost at the loose interfaces. Since the cryo-EM density for the Arg side chains in the Sensor II/Arginine finger motif are also weaker at the loose interfaces (Figure S1k), the GTP-binding sites at these locations are likely in a non-catalytic state. Taken together, these results indicate that TgMcrB^AAA^ possesses all the critical residues needed to bind and hydrolyze GTP.

### TgMcrB and TgMcrC form an asymmetric complex

We next sought to elucidate structural and biochemical consequences of TgMcrC binding to TgMcrB. Purified TgMcrC was very sensitive to buffer conditions and could only be concentrated in the presence of TgMcrB. Together full-length TgMcrB and TgMcrC formed stable, dumbbell-shaped complexes in the presence of GTPγS that were suitable for structure determination by cryo-EM (Figure S2a). Initial image processing showed that the complex consists of two TgMcrB hexamers connected through TgMcrC dimerization (Figure S2b). Because of structural variability, however, we were only able to refine a ‘half’-complex (Figure S2c), which yielded a map at an overall resolution of 2.4 Å (Figures 3 and S2d-g). In this reconstruction, a single TgMcrC binds the TgMcrB hexamer by inserting itself through the central pore of the AAA+ ring in an asymmetric fashion (Figure 3a-c).

**Figure 3.**
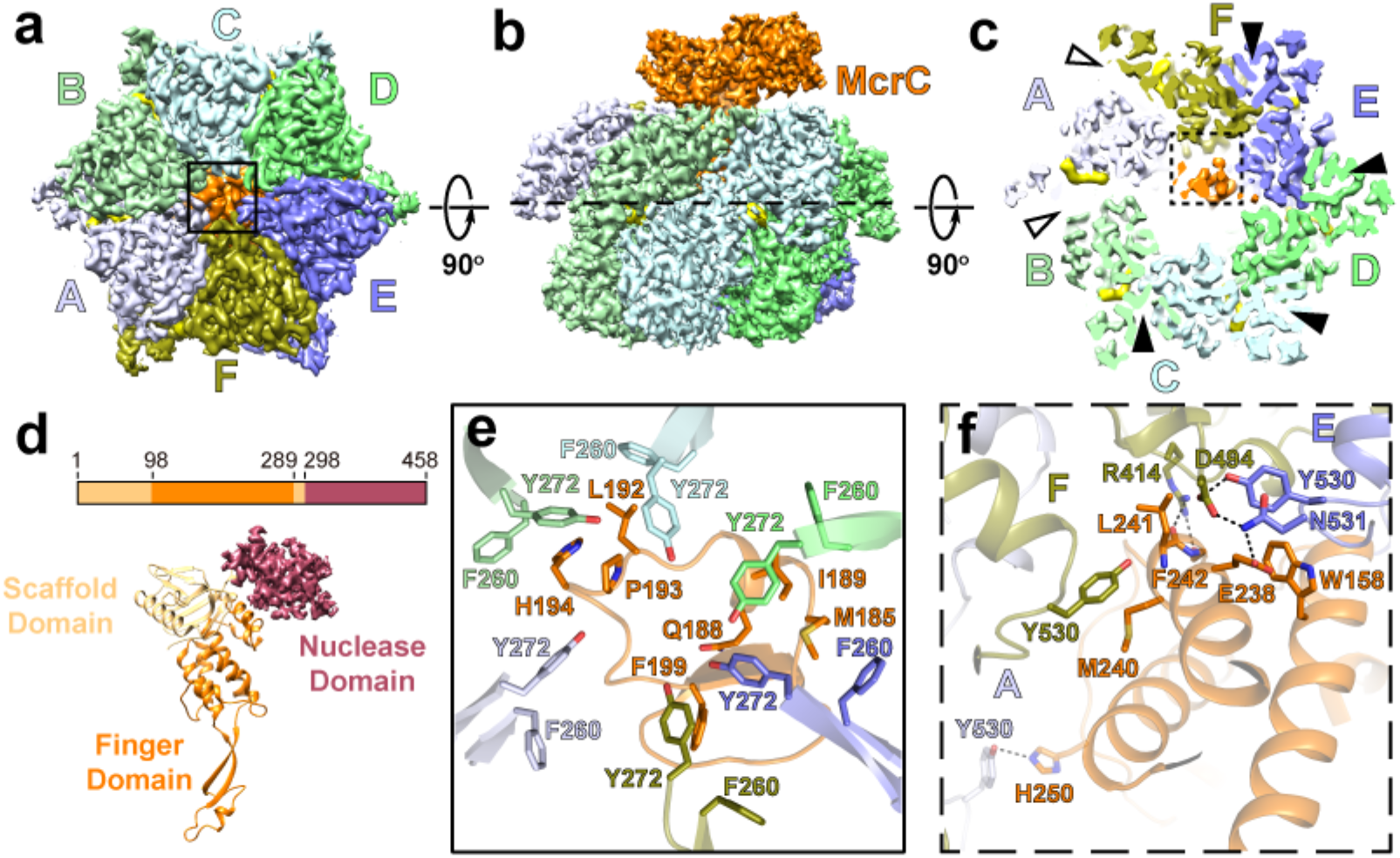
Asymmetric assembly of the TgMcrBC complex. (**a** and **b**) Bottom and side views of the cryo-EM density map of the TgMcrBC half-complex. TgMcrB subunits are colored as in Figure 1, TgMcrC is shown in orange, and nucleotides in yellow. (**c**) Slice section through the map of the TgMcrBC half-complex at the level of the bound nucleotides, indicated by the dashed line in (**b**). Solid and empty arrowheads indicate tight and loose interfaces, respectively. (**d**) Domain architecture of TgMcrC. (**e**) Close-up view of the interactions of TgMcrC with TgMcrB at the bottom of the hexamer, indicated by the black square in (**a**). (**f**) Close-up view of the interactions of TgMcrC with the TgMcrB hexamer at the E/F and F/A interfaces, indicated by the dashed black square in (**c**). Dashed lines indicate hydrogen bonds.

The resolution of our reconstruction allowed us to build the TgMcrC structure *de novo*. Each monomer contains a scaffold domain, a ‘finger domain’ and a C-terminal endonuclease domain (Figure 3d, Supplementary Data S3). The scaffold domain (residues 1-98 and 289-298) consists of a barrel-like structure that centrally positions the two flanking domains, forming a rigid connection between the finger and endonuclease domains. The finger domain (residues 99-288) adopts an extended, segmented structure with two antiparallel helices that contact the nuclease domain above, a helical bundle, and a long β-sheet ‘stalk’ that protrudes downward, terminating in a loop-helix-loop region at the tip (Figures 3d and S3a). The C-terminal endonuclease domain (residues 299-458) rests atop the structure and though poorly resolved in our map exhibits a fold characteristic of PD-(D/E)xK family enzymes.

The finger domain spans the entire length of the hexamer and its binding interface changes along the axis of the central pore (Figure S3a-e). At the top of the ring, the helical bundle associates with the F and E subunits and then tilts to contact the E and D subunits near the middle of the assembly (Figures 3c and S3a-c). We also observe interactions between the β-sheet stalk and the E subunit at this midpoint (Figure S3a and d). The loop-helix-loop at the distal tip of the finger domain plugs a narrow opening at the very bottom of the McrB hexamer (Figures 3a, S3a and e, Supplementary Data S3). Conserved aromatic residues Phe260 and Tyr272 from the helix 2 inserts of each McrB subunit surround and stabilize the tip (Figure 3e and Supplementary Data S1). While the finger domain interacts with all six subunits of TgMcrB at the bottom of the hexamer, TgMcrC binds the hexamer in a highly asymmetric fashion.

TgMcrC binding breaks the parallel π-stacking interaction between Arg414^F^ and Tyr530^E^ at the E/F interface (Figure 3c and f), which has the smallest interaction area among the four tight interfaces (~2400 Å^2^ versus >2500 Å^2^ for all the others). This perturbation changes the conformation of the 414-420 loop in subunit F as Arg414^F^ rotates to hydrogen bond with the main-chain atoms of Leu241^McrC^ and Phe242^McrC^ (Figures 3f and S3h-i). Concomitantly, Tyr530 and Asn531 in subunit E hydrogen bond to Asp494 in subunit F. Glu238 in the finger domain further stabilizes this conformation through an additional hydrogen bond with Asp494^F^. TgMcrC binding also generates some additional interactions in the F/A interface, where His250^McrC^ hydrogen bonds with Tyr530^McrB^ from the A subunit and Met240^McrC^ and Leu241^McrC^ form van der Waals interactions with Tyr530^McrB^ in the F subunit (Figure 3f, Supplementary Data S3). These interactions, which bury a combined surface area of 1298 Å^2^, serve to anchor McrC at the top of the ring, restricting its motion and orientation. Despite the localized differences at the E/F interface, the conformation of the TgMcrB hexamer remains largely unchanged in the TgMcrBC complex (overall RMSD of 0.75 Å compared to TgMcrB^AAA^ alone), with its intrinsic asymmetry and the remaining tight and loose interface interactions preserved (Figures 3c, S3f and g). These findings indicate that TgMcrC does not induce substantial remodeling of the TgMcrB hexamer but instead adapts and exploits its intrinsic asymmetry when binding.

### TgMcrC binding optimally positions existing catalytic machinery to stimulate GTP hydrolysis

A distinguishing feature of the *E. coli* McrBC system is the ability of McrC to stimulate McrB’s GTP hydrolysis *in vitro* (Pieper et al., 1997; Pieper et al., 1999a). Purified TgMcrC similarly stimulates TgMcrB’s basal GTPase activity, demonstrating that this is also a conserved property of other homologs (Figure 4d). Our high-resolution TgMcrBC structure reveals the underlying molecular mechanism governing this stimulation. As a consequence of the structural asymmetry imposed by the TgMcrB hexamer, TgMcrC’s finger domain engages only a single active site at a time (Figure 4a). Here, the helical bundle wedges against the D/E interface and inserts a highly conserved arginine (Arg263^McrC^) at the edge of the pocket (Figures 4b-c, S3a and c). Acting through a hydrogen-bonding network that includes Asn359^McrB^, Asn410^McrB^, Asp413^McrB^, and a bridging water (H_2_0^Bridging^), Arg263^McrC^ ultimately alters the conformation of the McrB consensus loop. This reorganization allows Asn410 and E357 of the Walker B motif to position a second water (H_2_0^Catalitic^) that is poised for nucleophilic attack on the γ-phosphate (Figure 4b). Glu357 also acts in concert with the Asp356 of the Walker B motif to stabilize a third water molecule that completes the octahedral coordination of the magnesium cofactor (Figure 4b).

**Figure 4.**
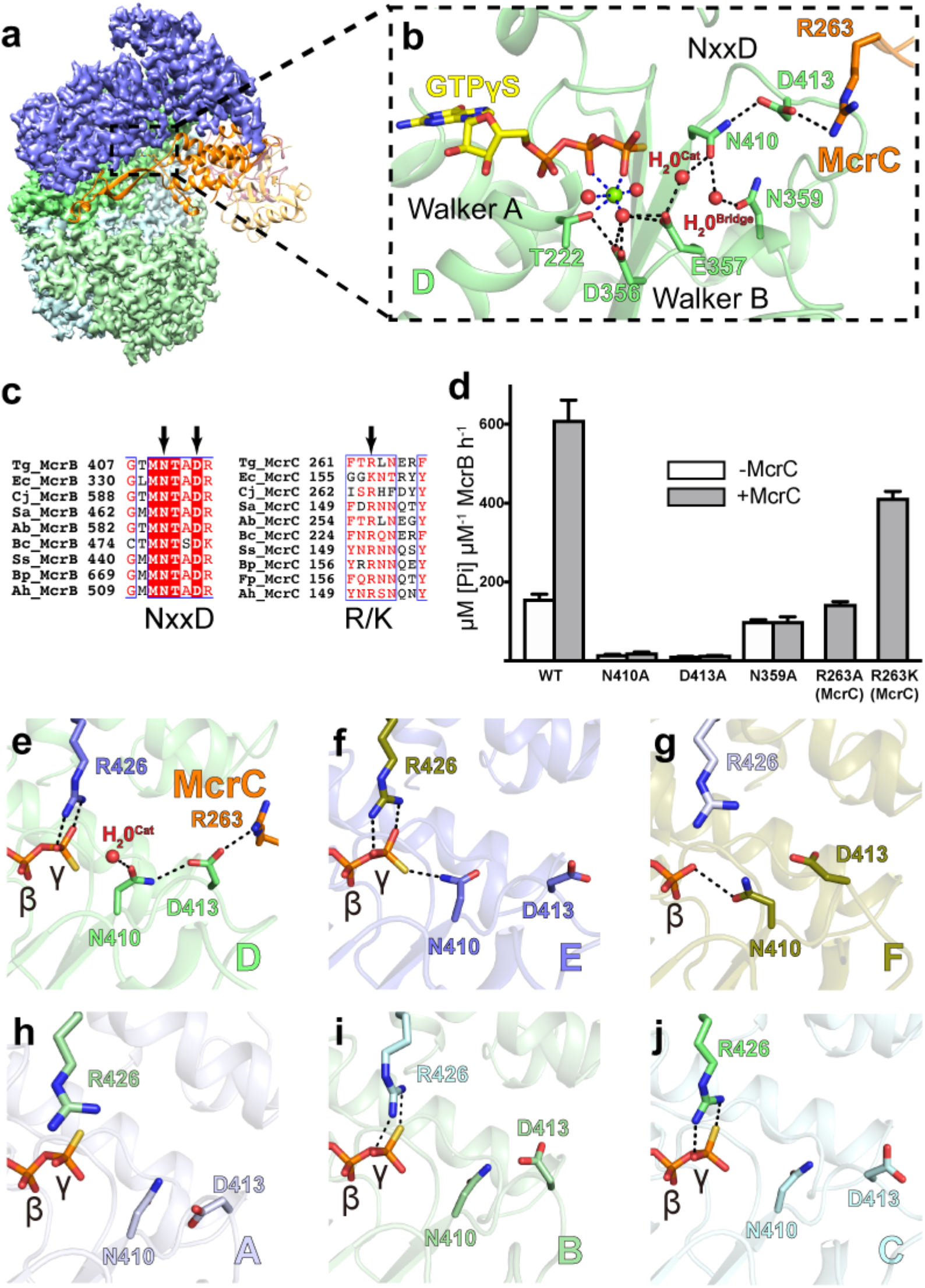
Structural basis for TgMcrC-mediated stimulation of TgMcrB GTPase activity. (**a**) Side view showing the interaction of TgMcrC with the D/E interface of the TgMcrB hexamer. TgMcrB and TgMcrC are colored as in Figure 3 and shown in surface and ribbon representation, respectively. For clarity, subunits A and F are not shown. (**b**) Hydrogen-bonding network formed by TgMcrC with residues of the NxxD motif at the D/E interface of the TgMcrB hexamer. Spheres indicate waters (red) and a magnesium ion (green). Dashed lines indicate hydrogen bonds (black) and metal coordination (blue). For clarity, the *trans* interacting residues in subunit E are not shown. (**c**) Sequence alignment of McrB and McrC homologs for the McrB signature sequence (NxxD) and the region in McrC that contains the inserted arginine/lysine residue (R/K). Abbreviations for the aligned species are as in Figure 2c. (**d**) Basal (-McrC) and TgMcrC-stimulated (+McrC) GTPase activity of TgMcrB for wild-type proteins and mutants with single amino-acid substitutions either of residues around the NxxD motif in TgMcrB or of residues in TgMcrC (n = 3, mean ± standard deviation). (**e–j**) Arrangement of the asparagine and aspartate residues of the NxxD motif at the six interfaces in the TgMcrB hexamer of the TgMcrBC complex.

Alanine substitutions at Asn410 and Asp413 in full-length TgMcrB abolish both basal and McrC-stimulated GTPase activity (Figure 4d), underscoring their crucial catalytic function. Mutation of Arg263^McrC^ to alanine selectively abrogates the stimulatory effect of McrC binding without impairing basal turnover (Figure 4d). The apparent GAP function thus arises from an indirect reconfiguration of the side chains that orient the catalytic water rather than promoting charge compensation in the transitions state.

### Sequential rearrangements of the consensus loop control the cycle of McrB GTP hydrolysis

The consensus loop and charge-compensating arginine finger (Arg426_*trans*_) adopt different conformations at each of the six interfaces within the McrC-bound TgMcrB hexamer (Figure 4e-j). As described above, the tight D/E interface shows an McrC-activated conformation with Arg426_*trans*_ stabilizing the γ-phosphate and Asn410 properly arranged to orient the catalytic water (Figure 4b and e). In the adjacent tight E/F interface, Asn410 and Arg426_*trans*_ appear in close contact with the γ-phosphate of GTPγS in a manner that excludes a potential catalytic water (Figure 4f). The loose F/A interface uniquely contains GDP with the side-chain oxygen of Asn410 forming a hydrogen bond with the β-phosphate. This partially occludes the space normally occupied by the γ-phosphate and forces Arg426_*trans*_ into a conformation in which it is angled away from the nucleotide (Figure 4g). At the loose A/B interface, Arg426_*trans*_ and Asn410 both point away from GTPγS, likely a consequence of the weakened inter-subunit interactions (Figure 4h). In both the tight B/C and C/D interfaces, Arg426_*trans*_ interacts with the γ-phosphate but Asn410 faces away from the nucleotide (Figure 4i and j). These pockets appear primed for hydrolysis but unable to proceed efficiently as Asp413 adopts random orientations in the absence of McrC and thus cannot help stably redirect Asn410 to position the catalytic water (Figure 4i and j).

These conformational differences likely reflect different states in the hydrolysis cycle, with the B/C and C/D active sites occupying a GTP-bound, pre-hydrolysis state, D/E most likely the activated transition state, E/F assuming a post-hydrolysis state, and the loose GDP-bound F/A and GTPγS-bound A/B sites depicting the phosphate release and subsequent nucleotide exchange steps, respectively. Together these data imply that TgMcrB GTP hydrolysis proceeds through a coordinated, sequential mechanism.

### McrBC homologs share a conserved architecture and catalytic mechanism

To establish whether different homologs use a conserved mechanism for stimulated hydrolysis, we determined the single-particle cryo-EM structure of the complex formed by the full-length *E. coli* proteins in the presence of GTPγS. EcMcrBC also formed dumbbell-shaped particles and we refined a ‘half’-map reconstruction of these assemblies to an overall resolution of 3.3 Å (Figures 5 and S4). As with TgMcrBC, a single EcMcrC monomer inserts into the central pore of the EcMcrB hexamer (Figure 5a and b). A cross-section slice through the map at the height of the bound nucleotides reveals that the same intrinsic asymmetry is present, with loose F/A and A/B interfaces and tight B/C, C/D, D/E and E/F interfaces (Figure 5c). The unique interactions stabilizing each tight interface are also conserved in EcMcrBC and absent in the loose interfaces: Arg337 (Arg414 in TgMcrB) and Asp343 (Asp420 in TgMcrB) interact *in trans* with Phe428 (Tyr530 in TgMcrB) and Arg283 (Arg360 in TgMcrB), respectively (Figure S5a and b). In contrast to the TgMcrBC structure, we observe unambiguous density for GDP in the A and F monomers of the EcMcrB hexamer (Figure 5c).

**Figure 5.**
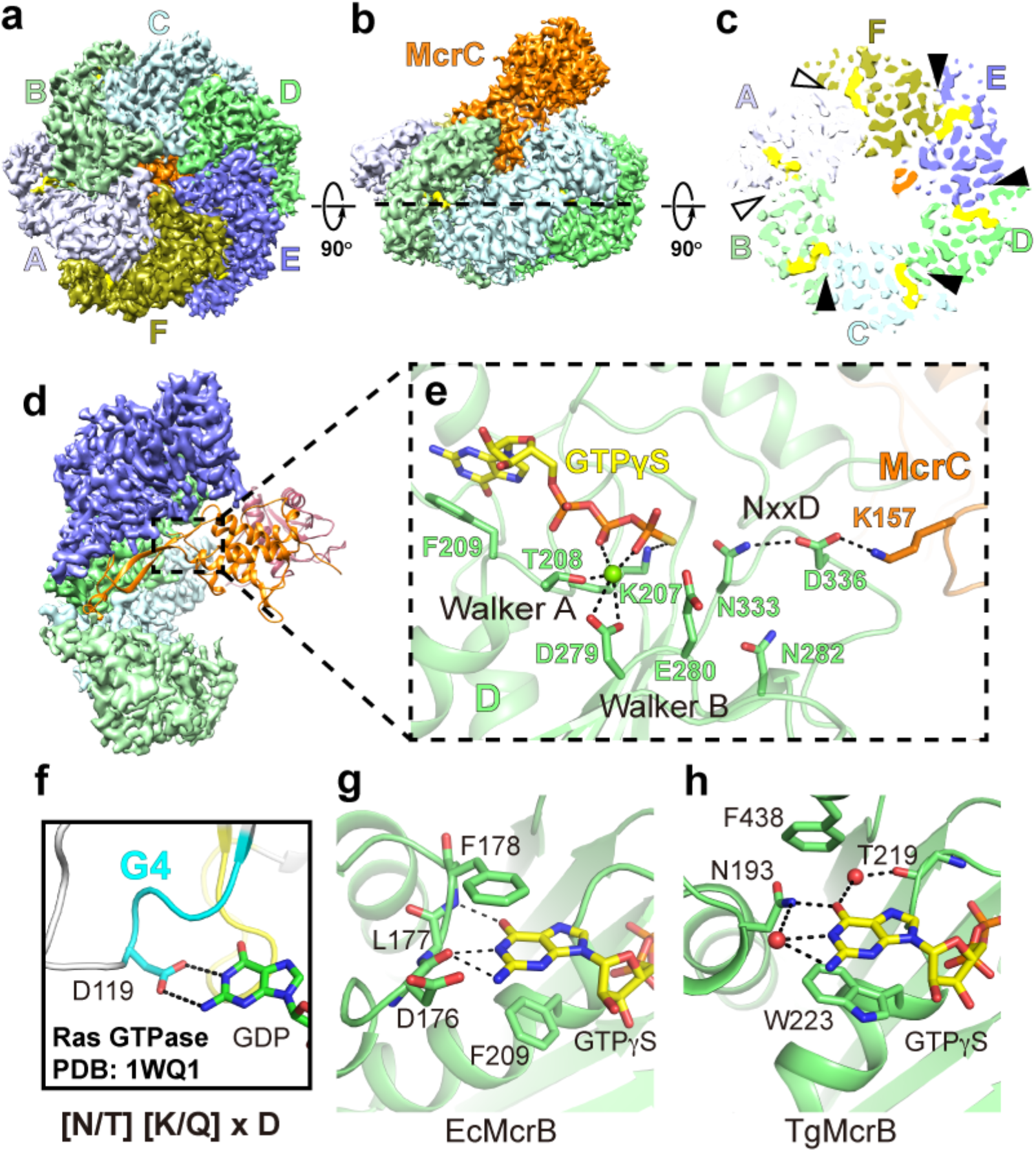
The McrBC complexes of *E. coli* and *T. gammatolerans* show a conserved architecture. (**a** and **b**) Bottom and side views of the cryo-EM density map of the EcMcrBC half-complex. (**c**) Slice section through the map of the EcMcrBC half-complex at the level of the bound nucleotides, indicated by the dashed line in (**b**). Solid and empty arrowheads indicate tight and loose interfaces, respectively. (**d**) Close-up view of the interaction of EcMcrC with the EcMcrB hexamer at the D/E interface. EcMcrB and EcMcrC are shown in surface and ribbon representation, respectively. For clarity, subunits A and F are not shown. (**e**) Hydrogen-bonding network formed by EcMcrC with residues of the NxxD motif at the D/E interface of the EcMcrB hexamer. Spheres indicate waters (red) and a magnesium ion (green). Dashed lines indicate hydrogen bonds and metal coordination. (**f-h**) Structural basis for guanine recognition in the Ras GTPase (PDB: 1WQ1; Scheffzek et al., 1997) and in the McrB homologs (EcMcrB and TgMcrB).

EcMcrC shares the same general architecture as TgMcrC, featuring an extended finger domain and a C-terminal nuclease domain. EcMcrC, however, lacks the N-terminal portion of the scaffold domain, retaining only a small β-hairpin insertion between the finger and nuclease domains (Figures 3d and S5e). Sequence alignment of McrC family proteins suggests that these insertion strands serve as a conserved linker between the finger and nuclease domains (Supplementary Data S3). The finger domains superimpose with an RMSD of 2.4 Å, confirming the overall structural conservation between these evolutionarily remote homologs.

The structural asymmetry present in the EcMcrBC complex similarly biases EcMcrC to associate with only a single active site at a time (Figure 5c and d). EcMcrC inserts Lys157 into the D/E interface of the EcMcrB hexamer and employs the same hydrogen-bonding network seen in the TgMcrBC complex to reorient Asn333 and asp336 in the McrB signature motif (Figure 5e). Asn282 spatially occupies the same position as Asn359 in TgMcrB (Figures 4b and 5e). Although we do not resolve the catalytic or bridging waters in our structure of the *E. coli* complex, the location of this side chain suggests a conserved mechanistic function. The rest of the catalytic machinery is also conserved (Figures 5e, S5c and d).

Our structural findings rationalize previous phenotypes associated with consensus loop mutants in EcMcrB. Asn333Ala and Asp336Asn substitutions would disrupt the hydrogen-bonding network needed to position the catalytic water, leading to a complete loss of GTPase activity and the abrogation of long-range DNA cleavage (Pieper et al., 1997; Pieper et al., 1999a). Loss of stimulated GTPase activity due to an alanine mutation at Asn282 would arise from a similar structural perturbation (Pieper et al., 1999a). Interestingly, substituting a lysine for TgMcrC’s catalytic Arg263 partially restores the stimulatory effect that is lost when this side chain is replaced with alanine (Figure 4d). Together these data demonstrate that stimulated GTP hydrolysis in different McrBC homologs occurs via a conserved molecular mechanism.

### Divergent McrB homologs employ the same generalized principles for nucleotide specificity

In every GTPase, the conserved sequence [N/T] [K/Q]xD (termed the ‘G4 element’) confers nucleotide specificity (Bourne et al., 1991; Leipe et al., 2002; Paduch et al., 2001). The absolutely conserved aspartate side chain in this motif forms specific hydrogen bonds with the 1’ amine and 2’ amino group of the guanine base, thereby distinguishing it from ATP (Figure 5f). Nothing in the TgMcrB AAA+ domain makes contact with this portion of the nucleotide (Figure 2a and b), suggesting other structural features fulfill this role. Our reconstructions of the full-length EcMcrBC and TgMcrBC complexes reveal how each individually achieves this end (Figure 5g and h). In EcMcrBC, a loop that lies directly upstream of the AAA+ domain coordinates the guanine base through main-chain interactions (Figure 5g). The backbone carbonyl of Leu177 hydrogen bonds with both the 1’ amine and 2’ amino group of the guanine base, while the main chain nitrogen of Phe178 reads out the 6’ carbonyl group. Collectively these interactions would discriminate against the substitution of an amino group at the 6’ position (as in ATP and XTP) and the absence of an amino group at the 2’ position (as in ATP and ITP), consistent with EcMcrB’s nucleotide selectivity preferences of GTP>ITP >XTP>>ATP (Pieper et al., 1999a). TgMcrB, in contrast, specifically coordinates the guanine base through two water-mediated interactions (Figure 5h). Asn193 at the very beginning of the AAA+ domain directly hydrogen bonds to guanine’s 6’ carbonyl and orients a water molecule to interact with the 1’ amine and 2’ amino group. The backbone carbonyl of Thr219 also interacts with the 6’ carbonyl group of the base via a second bridging water. Importantly, the fundamental chemistry underlying guanine nucleotide recognition is conserved between both homologs despite each utilizing different structural elements.

### McrBC forms a tetradecameric assembly through the dimerization of McrC

Previous studies reported that EcMcrBC complexes form tetradecameric assemblies *in vitro* (Nirwan et al., 2019a; Panne et al., 2001). In our hands, dimeric McrBC complexes generated using the full-length Tg and Ec proteins exhibit a high degree of conformational variability, which prevented us from calculating interpretable maps for these larger oligomeric states and limited our ability to analyze the dimer interface between the two McrC subunits. To overcome this limitation, we produced complexes containing full-length TgMcrC bound to the AAA+ domain of TgMcrB (TgMcrB^AAA^C) in the presence of GTPγS. This assembly was structurally more homogeneous and allowed us to calculate maps of the ‘half’-complex’ at 3.7-Å resolution as well as a C2-symmetrized map of the entire TgMcrB^AAA^C tetradecameric complex at 4.2-Å resolution (Figure S6). A TgMcrC dimer bridges two TgMcrB^AAA^ hexamers in this structure (Figure 6a), with the scaffold and nuclease domains forming the dimer interface (Figure 6b). The nuclease domains associate through their α12 helices and a loop between the β10 and β11 strands (‘L’), whereas the neighboring scaffold domains interact with each other through their β4 strands that form main-chain hydrogen bonds with each other. While we could not calculate a reconstruction for the full EcMcrBC tetradecameric assembly, the map for the half-complex contains density for an additional ordered nuclease domain of EcMcrC (Figure S5f). The organization of the EcMcrC nuclease domains at the dimer interface is identical to that seen in the TgMcrC dimer, with the α10 helix and an analogous extended loop serving as the primary points of contact (Figure S7a).

**Figure 6.**
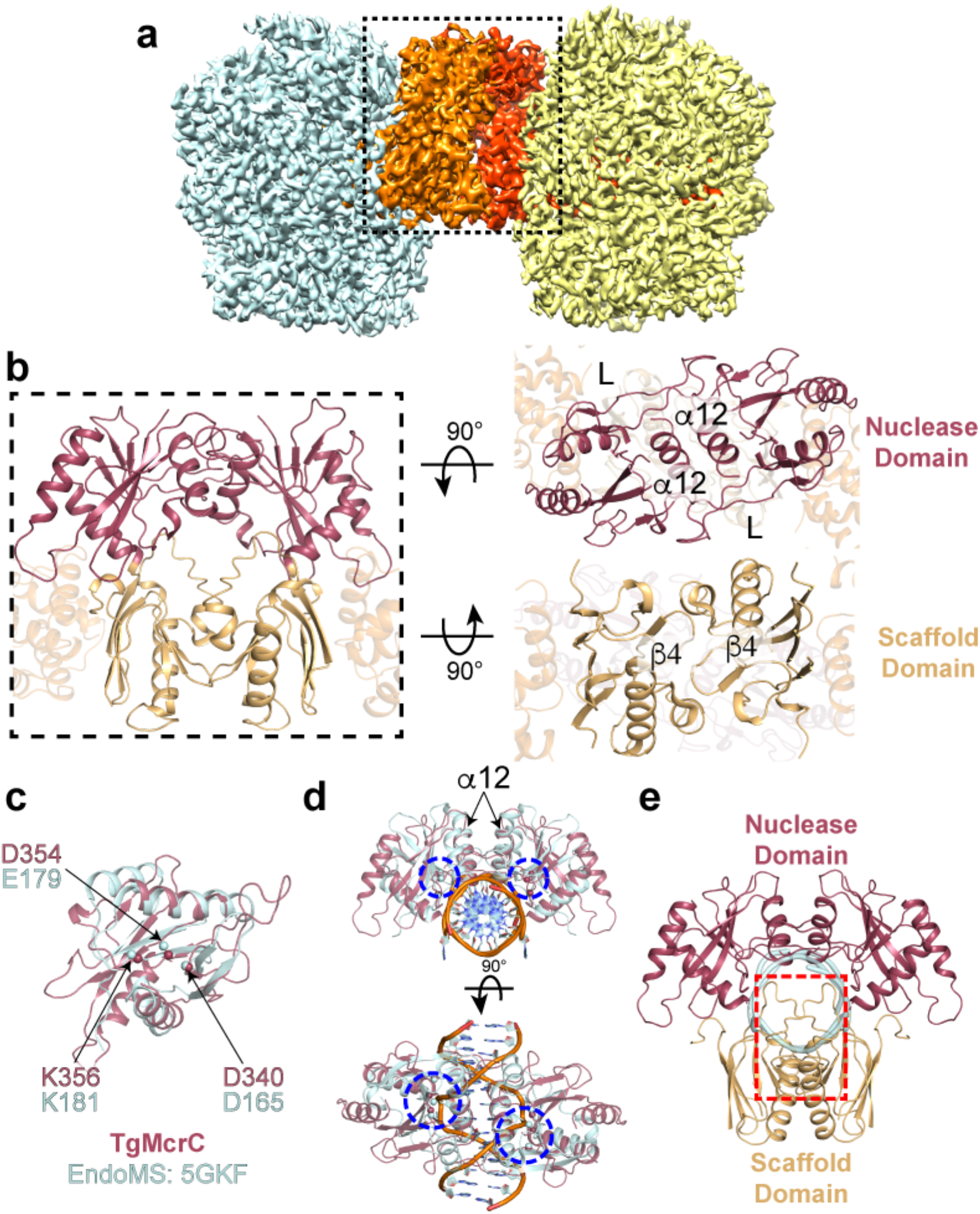
The tetradecameric assembly of the TgMcrB^AAA^C complex shows a cleavage-incompetent conformation. (**a**) Composite cryo-EM map of the full TgMcrB^AAA^C complex. Two TgMcrC (orange and red) form a dimer that bridges two TgMcrB hexamers (cyan and yellow). (**b**) Close-up views of the TgMcrC dimer interfaces formed by the two nuclease domains (upper right panel) and the two N-terminal domains (lower right panel). (**c**) Superposition of the monomeric structures of TgMcrC and EndoMS (PDB: 5GKF; Nakae et al., 2016). The conserved residues involved in the cleavage activity are labeled and shown as spheres. (**d**) Structural comparison between the TgMcrC dimer in the TgMcrB^AAA^C complex and the EndoMS dimer in a DNA-bound state. The blue dashed circles indicate the active sites for DNA cleavage. (**e**) Illustration of the cleavage-incompetent conformation of TgMcrC. For clarity, the structure of the EndoMS protein is not shown. The backbone of the DNA substrate bound to EndoMS is colored in cyan. The red square indicates the regions of potential steric clashes.

### The McrC dimer adopts a cleavage-incompetent conformation in the absence of a DNA substrate

The DNA-bound structures of other PD-(D/E)xK nucleases provide a template for modeling McrC’s cleavage activity. Of the many structural homologs identified by the DALI server (Holm and Rosenstrom, 2010), the coordinates of the *Thermococcus kodakarensis* EndoMS endonuclease (PDB: 5GKF; Z-score 7.9; Nakae et al., 2016) provided the best framework for these purposes. EndoMS binds DNA as a dimer, with each active site attacking a single strand of the DNA duplex to induce a double-strand break. As with other PD-(D/E)xK enzymes, Asp165^EndoMS^, Glu179^EndoMS^ and Lys181^EndoMS^ coordinate a divalent metal cofactor that is required for catalytic function (Knizewski et al., 2007; Nakae et al., 2016). Structural superposition confirms TgMcrC’s C terminus shares the same fold and identifies Asp340^TgMcrC^, Asp354^TgMcrC^ and Lys356^TgMcrC^ as putative catalytic side chains based on their spatial alignment with the EndoMS metal-binding residues (Figure 6c). EcMcrC also shares this structural homology (Figure S7b). Importantly, our modeling is consistent with previous biochemical data showing that mutation of the predicted catalytic residues in EcMcrC (Asp224^EcMcrC^, Asp257^EcMcrC^ and Lys259^EcMcC^) impairs cleavage of modified DNA *in vitro* (Pieper and Pingoud, 2002). Further comparison shows that the organization and location of the active sites in the TgMcrC and EcMcrC dimers is conserved between the two species (Figure S7c).

To gain insight into McrC cleavage, we overlaid two copies of the TgMcrC and EcMcrC endonuclease domains independently onto the dimeric, DNA-bound EndoMS complex (Figures 6d and S7d). The nuclease domains align in an orientation that resembles the dimer configuration captured in our cryo-EM structures; however, we observe numerous steric clashes in both models. TgMcrC’s scaffold domain and the α12 nuclease helices collide with the DNA substrate (Figure 6e). EcMcrC lacks an N-terminal scaffold domain yet still clashes with the DNA backbone, owing to the first helix of its nuclease domain being significantly longer (Figure S7e). Attempts to model similar interactions with other structurally related homologs like EcoRV (PDB: 1AZ0; Perona and Martin, 1997) and the *Sulfolobus solfataricus* Holliday junction endonuclease (PDB: 1OB8; Middleton et al., 2004) resulted in substantial clashes between the two McrB hexamers. We therefore speculate that our dimeric McrBC structures depict a conformation that is incompatible with DNA cleavage and that a major conformational change would be required for nuclease activity to proceed unencumbered.

## DISCUSSION

Our structural analysis reveals that TgMcrB^AAA^ forms an asymmetric hexamer, similar to the architecture adopted by many other AAA+ family proteins (de la Pena et al., 2018; Enemark and Joshua-Tor, 2006; Gates et al., 2017; Puchades et al., 2017; Ripstein et al., 2017; Twomey et al., 2019; Zehr et al., 2017; Zhao et al., 2015). Asymmetry appears to be maintained by the conformation of key interface residues – Arg360, Glu527 and Tyr530 in one monomer and Arg414, Asp420 and Arg424 in its neighbor – acting *in trans*. Alanine substitutions of these residues increase basal GTPase activity by ~two-fold (Figure S1i), suggesting they help restrict uncoordinated, random GTP hydrolysis throughout the hexamer. The asymmetry in the ring also explains how crystal packing forces could distort the hexamer at the two loose interfaces leading to the open ring conformation observed in our TgMcrB^AAA^ X-ray structure (Figure S1j). These observations argue that asymmetry is an intrinsic characteristic of the McrB^AAA^ hexamer rather than being induced upon McrC binding as has recently been proposed (Nirwan et al., 2019b).

The TgMcrB AAA+ domain possesses all the catalytic machinery needed for nucleotide hydrolysis. We find that the canonical *cis*-acting Sensor II arginine is replaced with a *trans*-acting arginine (Arg425) that is positioned adjacent to the charge-compensating arginine finger (Arg426) in helix α11 (Figure 2b). Our cryo-EM and X-ray structures of TgMcrB^AAA^ reveal that Arg425 is not only important for stabilizing Glu375 *in cis* as predicted from the previous structures of *E. coli* complexes (Nirwan et al., 2019b) but also interacts with the phosphates of GTP *in trans* (Figure 2b). Asn410 (consensus loop) and Glu357 (Walker B motif) together position the catalytic water. We note that Trp223 forms a crucial π-stacking interaction with the guanine base that is also present in EcMcrB and functionally conserved at the sequence level in other homologs. Perturbing any of these side chains reduces basal GTP hydrolysis of TgMcrB^AAA^. Similar phenotypes were observed with the corresponding mutations in the *E. coli* protein (Pieper et al., 1999a), indicating that the basic catalytic machinery is hardwired into the McrB AAA+ fold across evolution.

We demonstrate that McrC-stimulated GTP hydrolysis is a broadly conserved property of the McrBC family and not simply a unique feature of the *E. coli* homolog (Figure 4d) (Pieper et al., 1997; Pieper et al., 1999a). While this type of stimulation is uncommon among AAA+ proteins, it resembles the activation of small G proteins by their cognate GAPs. GAPs enhance catalytic turnover either by contributing essential catalytic residues *in trans* or by conformationally stabilizing and/or reorienting active-site elements into an optimal configuration (Bos et al., 2007). In nearly every case, these interactions affect the charge-compensating element (Mishra and Lambright, 2016). RasGAP, for example, provides the arginine finger needed for Ras turnover while RGS4 binding to G_*iα1*_ reorients an existing arginine in the switch I motif (Scheffzek et al., 1997; Tesmer et al., 1997). A notable exception is RapGAP, which provides *in trans* an asparagine that positions the nucleophilic water (Scrima et al., 2008). Our structures show that TgMcrC and EcMcrC stimulate hydrolysis indirectly by altering the conformation of the McrB consensus loop. Both proteins insert a conserved basic residue (Arg263^TgMcrC^ and Lys157^EcMCrC^) at the edge of the McrB active site (Figure 4c) and, via a hydrogen-bonding network, reposition a conserved asparagine (Asn410^TgMcrB^ and Asn333^EcMcrB^) that in turn correctly orients the catalytic water for nucleophilic attack on the γ-phosphate (Figures 4b and 5e). This conserved molecular mechanism thus represents a unique variation on a common theme. We note that the helical bundle of the McrC finger domain wedges into the E/F interface at the top of the McrB hexamer in both structures (Figure 3f). This interaction not only anchors McrC but also directs its catalytic machinery to the adjacent active site at the D/E interface (Figures 4b and S3c). These constraints dictate that McrC stimulation can only occur at a single active site at any given time.

In our structures, the consensus loop and the *in trans* arginine finger adopt different conformations in each active site around the McrB hexamer (Figures 4e-j). We interpret each configuration as representing a different state in the hydrolysis cycle (Figure 7, dashed red outline). The McrC-engaged D/E active site assumes a transition state-like conformation with the catalytic machinery optimally positioned for simulated turnover. In the tight C/D and B/C active sites, GTP is bound but the catalytic components are in a suboptimal conformation. This configuration suggests a pre-hydrolysis state that is primed for interaction with McrC. The loose A/B and F/A sites represent low-affinity, post-hydrolysis states that allow for free exchange of GTP and GDP, consistent with McrB not requiring a guanine nucleotide exchange factor. GDP occupies both sites in our EcMcrBC structure (Figure 5c) while we observe GTPγS in the A/B site of the TgMcrBC complex (Figure 1c), indicating nucleotide exchange has already occurred. The final tight E/F site likely adopts a post-hydrolysis state that is partially destabilized but still remains intact due to the presence of the γ-phosphate. These data suggest that McrC-stimulated GTP hydrolysis proceeds via a coordinated mechanism that cycles around the McrB hexamer, engaging each composite active site sequentially (Figure 7). In this scheme, the release of the γ-phosphate and the intrinsic asymmetry of the complex serve as the driving forces for a rotational movement. Release of the phosphate would destabilize the E/F interface, converting it from a tight to a loose configuration. This could promote a transition of the A/B interface from loose to tight, where exchange of GTP for GDP has presumably occurred. Weakening the E/F interface would destabilize the interactions with the helical bundle that anchor the finger domain (Figure 3f), thereby releasing McrC and allowing it to rotate. The asymmetry of the structure would bias the movement in a clockwise direction as the helical bundle of the finger domain would not be able to associate with the loose F/A interface and thus would have to intercalate into the D/E interface. This engagement would orient McrC to insert its catalytic arginine/lysine into the C/D active site, where it could trigger the next hydrolysis event to power the motor (Figure 7). The extensive contacts formed between the helix-loop-helix tip of the finger domain and all six subunits of the McrB hexamer (Figure 3e) would ensure that McrC does not dissociate from the complex following stimulated turnover. The stepwise transition from one binding interface to the next (Figure 7) is reminiscent of F/V-type ATPases (Forgac, 2007; Iwata et al., 2004; Stock et al., 2000; Yasuda et al., 2001). Given the conserved structural features and asymmetry present in both the Tg and Ec complexes, we anticipate that other McrBC homologs will follow this mechanochemical model.

**Figure 7:**
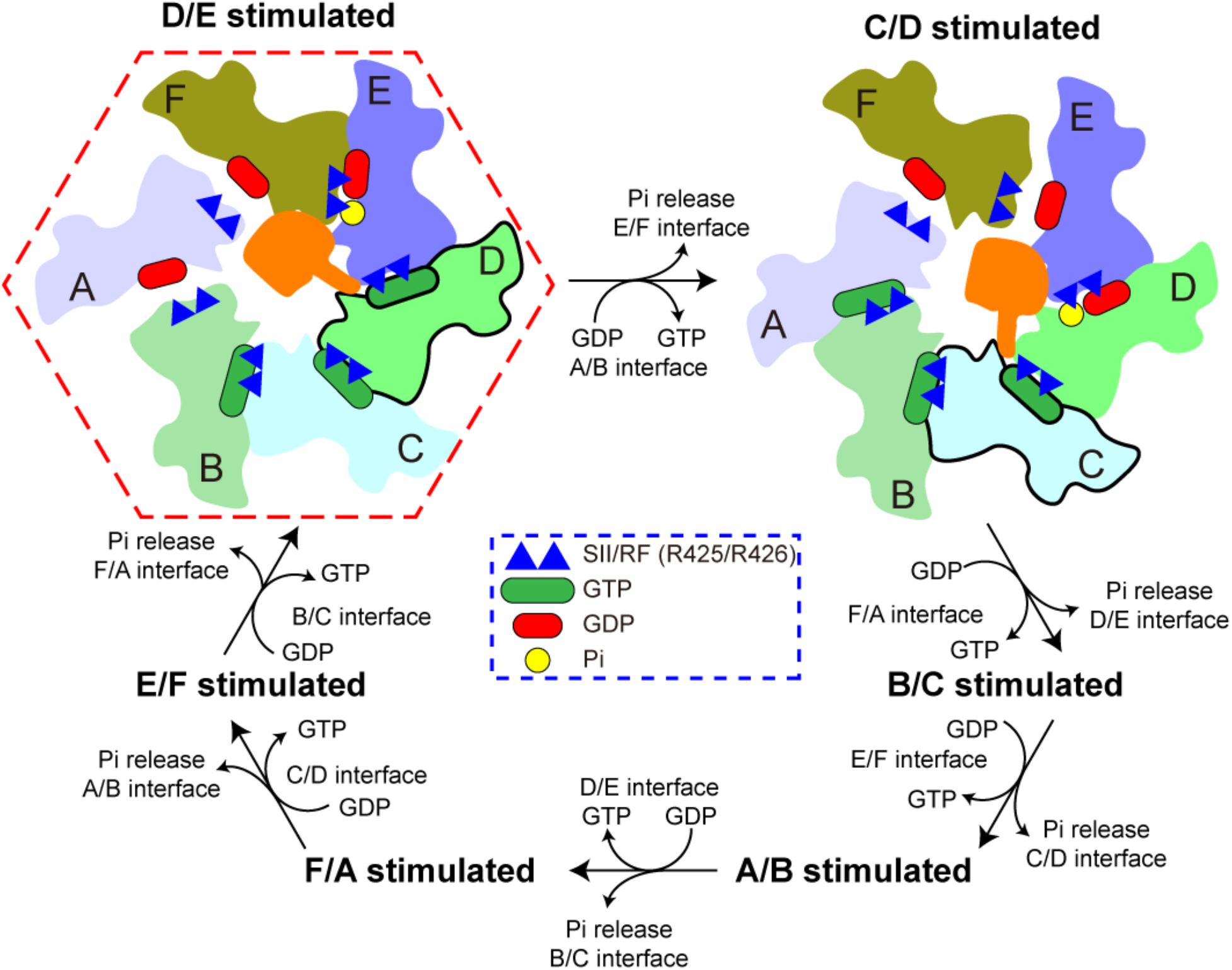
Rotation model for the catalytic cycle of McrB. Schematic representation of the putative GTP hydrolysis cycle that proceeds sequentially in a clockwise manner around the hexameric McrB ring relative to McrC in the central pore. The ‘finger’ extending from McrC represents the arginine/lysine residue that interacts with the NxxD motif. Ovals at the interfaces of the hexamer represent GTP (green) and GDP (red). The subunits indicated by the thick outlines are those in which the NxxD has been reorganized by McrC inserting its arginine/lysine residue. The red hexagon indicates the state observed in our cryo-EM structures.

Efficient hydrolysis also depends on an enzyme’s ability to bind and differentiate its appropriate nucleotide substrate. GTPases use the conserved aspartate in the G4 element to coordinate substituents at the 1’ and 2’ positions of the guanine base while AAA+ proteins recognize the amino group at the 6’ position in adenine (Bourne et al., 1991; Scheffzek et al., 1997). By reading out the 1’, 2’, and 6’ positions of the guanine base, McrB homologs appear to have combined both strategies to fine-tune their specificity for GTP in the context of a AAA+ fold. Ec and TgMcrB both use the same basic chemistry for this recognition, but each employs different structural components to mediate these contacts (Figure 5g-h). Interestingly, these pieces lie outside the core AAA+ fold and localize to either the flexible linker that connects to EcMcrB’s N-terminal DNA-binding domain and or the very start of helix α1 in TgMcrB (Supplementary Data S1 and S2a, colored in gold). Although the motor and cleavage machineries are conserved among McrB homologs (Figures 3-5), the N-terminal domains and connecting linkers are highly divergent. Crystallographic studies have shown that EcMcrB uses the DUF3578 fold to bind methyl-cytosine modifications (Sukackaite et al., 2012; Zagorskaite et al., 2018), whereas the N-terminal domain of TgMcrB consists of a YTH fold that specifically targets 6mA-modified DNA (Hosford et al., 2020). The related LlaJI restriction system from *Helicobacter pylori* binds DNA site-specifically via an N-terminal B3 domain (Hosford and Chappie, 2018). The subtle distinctions we observe with regard to nucleotide recognition are therefore significant and provide a blueprint for how divergent homologs can adapt the same fundamental chemistry to radically different structural contexts. Future structural characterization will determine if these principles hold true for other McrBC family members.

Previous biochemical studies suggest that McrBC’s stimulated GTP hydrolysis powers DNA translocation (Panne et al., 1999; Sutherland et al., 1992). While we do not directly address how this may occur in this study, our structures impose constraints with regard to the potential pathway of DNA and the organization of a cleavage-competent McrBC complex. DNA and RNA typically pass through the central pore of hexameric AAA+ helicases and translocases driven by ATP hydrolysis (Enemark and Joshua-Tor, 2006; Meagher et al., 2019). Based on recent cryo-EM reconstructions, a similar mechanism has been proposed for EcMrBC, in which the McrB N-terminal domains might interact with DNA on the bottom of the hexamer and thread it into the central channel (Nirwan et al., 2019b). Although we see in our map of full-length EcMcrBC weak density corresponding to the N-terminal domains near the top of the complex (Figure S7f), numerous structural observations oppose this potential trajectory. First, McrC specifically binds in the center of the McrB hexamer, blocking access to this pathway in both the Tg and Ec complexes. The asymmetric association of the finger domain’s helical bundle with the D/E/F subunits shrinks the pore diameter at the top of the hexamer from ~50 Å to ~15 Å (Figure S3b) while the loop-helix-loop region completely occludes the pore at the bottom of the hexamer (Figures 3a, S3e and S5a). Passage through the ring in this state would require both distortion and/or melting of the DNA duplex to conform to the narrow dimensions of the structure as well as either a complete displacement or gross conformational reorganization of McrC. Such changes would uncouple the sequential, coordinated stimulation of GTP hydrolysis suggested by our structures and yield a translocation mechanism that would use a completely stochastic catalytic process and would depend on alternating cycles of binding and dissociation for both McrC and DNA. While we cannot rule out that additional conformational changes occur upon DNA binding, biochemical characterization of EcMcrBC has shown that DNA binding and GTP hydrolysis are separate and distinct properties *in vitro* (Gast et al., 1997; Panne et al., 1999; Pieper et al., 1997; Pieper et al., 1999b). It therefore seems unlikely that DNA binding would significantly alter the architectural and catalytic interactions that have been conserved across kingdoms. Second, we resolve clear density decorating the outside edges of the TgMcrB^AAA^ hexamer that we attribute to the TgMcrB N-terminal domains (Figure S7g). The localization of these domains nearly perpendicular to the pore axis would require DNA, if it were to pass through the center of the TgMcrBC complex, to bend dramatically, more than has been observed in any structure to date. Energetically, such a configuration would be extremely unfavorable (Peters and Maher, 2010). The short seven amino acid linker connecting the N-terminal domains to the Tg AAA+ domains combined with the structural requirements of nucleotide selectivity would likely prohibit a large-scale rearrangement of these domains within the restriction complex. Taken together, these findings argue against a mechanism in which DNA passes through the central channel in the McrB hexamer. We speculate that McrBC complexes use a novel, yet to be elucidated means to translocate DNA.

Both EcMcrBC and TgMcrBC form tetradecameric complexes that are bridged by an McrC dimer (Figures S4 and S6). Our structural modeling, however, suggests that the conformation of this McrC dimer is incompatible with DNA binding and cleavage. Superposition with EndoMS shows that the N-terminal scaffold domain of TgMcrC and the first helix in the nuclease domain of EcMcrC clash with the modeled DNA substrate (Figures 6d and e, S7d and e). Modeling with other structurally related homologs produced more extreme clashing between the two McrB hexamers. It remains to be seen whether DNA binding alone could induce a cleavage-competent conformation. Interestingly, GTPγS does not support EcMcrBC DNA cleavage *in vitro* (Panne et al., 1999), consistent with our structural findings here. Moreover, mutation of Pro203 to valine in EcMcrB significantly reduces both EcMcrC-stimulated GTP hydrolysis and DNA cleavage of an ‘ideal’ substrate with R^M^C sites optimally spaced 63 base pairs apart so as not to require translocation (Panne et al., 1999). This finding raises the possibility that GTP hydrolysis is also needed for the transient reorganization of the McrC monomers, and that blocking this activity would lead to a non-productive arrangement. Further experiments will be needed to fully understand how the McrBC complex cleaves DNA.

Modification-dependent restriction systems function as a conserved barrier to lytic phage infections. In the ongoing arms race between virus and host, phages have evolved inhibitors against McrBC and GmrSD (Bair and Black, 2007; Dharmalingam and Goldberg, 1976), which confer the ability to bypass these defense machineries and allow phages to survive under conditions in which they would normally be restricted. Knowing how these defense systems work and how they have been naturally subverted is clinically important and will aid in the long-term development of small-molecule inhibitors that can impair conserved defense systems and improve the efficacy of phage-based treatments.

## ACKNOWLEDGEMENTS

We are grateful to Mark Ebrahim and Johanna Sotiris for help with grid screening and data collection at the Evelyn Gruss Lipper Cryo-Electron Microscopy Resource Center at The Rockefeller University. We thank Drs. Richard Cerione and Holger Sondermann for critical reading of the manuscript and the Northeastern Collaborative Access Team (NE-CAT) beamline staff at the Advanced Photon Source (APS) for assistance with remote X-ray data collection. This work was supported by National Institutes of Health Grant GM120242 (to J.S.C.) and is based upon research conducted at NE-CAT beamlines (24-ID-C and 24-ID-E) under the general user proposals GUP-51113 and GUP-41829 (PI: J.S.C.). NE-CAT beamlines are funded by the National Institute of General Medical Sciences from the National Institutes of Health (P30 GM124165). The Pilatus 6M detector on 24-ID-C beam line is funded by a NIH-ORIP HEI grant (S10 RR029205). This research used resources of the Advanced Photon Source, a U.S. Department of Energy (DOE) Office of Science User Facility operated for the DOE Office of Science by Argonne National Laboratory under Contract No. DE-AC02-06CH11357. J.S.C. is a Meinig Family Investigator in the Life Sciences.

## Author Contributions

Y.N., H.S., C.J.H., T.W., and J.S.C. designed the study and analysed data. C.J.H. and Y.N. cloned purified all constructs and carried out biochemical assays. Y.N. and H.S. collected cryo-EM data, carried out image processing, and built the atomic-resolution cryo-EM models. C.J.H. and Y.N. collected X-ray diffraction data, determined the X-ray structure, and built the X-ray model. Y.N., H.S., T.W., and J.S.C. wrote the manuscript.

