## Supplementary Information for "Structural asymmetry governs the assembly and GTPase activity of McrBC restriction complexes"

**Methods**

**Supplementary References**

**Supplementary Data**

Supplementary Data S1. Sequence alignment of the McrB family proteins.

Supplementary Data S2. Secondary structure topology diagram and monomeric structure of the AAA+ domains of TgMcrB and EcMcrB in ribbon representations.

Supplementary Data S3. Sequence alignment of the McrC family proteins.

**Supplementary Figures**

Figure S1: Biochemical and structural analysis of the TgMcrB$^{AAA}$ hexamer, related to Figures 1 and 2.

Figure S2: Cryo-EM analysis of the TgMcrBC complex, related to Figures 3 and 4.

Figure S3: Bound TgMcrC adopts to the intrinsic asymmetry of the TgMcrB hexamer, related to Figure 3 and 4.

Figure S4: Cryo-EM analysis of the EcMcrBC complex, related to Figure 5.

Figure S5: Cryo-EM structure of the EcMcrBC complex, related to Figure 5.

Figure S6: Cryo-EM analysis of the TgMcrB$^{AAA}$C complex, related to Figure 6.

Figure S7: Structural comparison of the nuclease domains, related to Figure 6.

**Supplementary Tables**

Supplementary Table S1. X-ray data collection and refinement statistics.

Supplementary Table S2. Cryo-EM data collection and refinement statistics.

### Methods

#### Cloning, expression and purification of TgMcrB$^{AAA}$

The gene for the *T. gammatolerans EJ3* McrB protein (DOE IMG/M ID 644807740) was codon optimized for expression in *E. coli* and synthesized commercially by GENEART. The DNA for the AAA domain of TgMcrB (residues 186-613) was amplified by PCR and cloned into the pET15bp vector, a modified version of the pET15b vector, in which the Factor Xa cleavage site after the N-terminal 6xHis tag was replaced with an HRV 3C cleavage site. Gibson assembly (New England Biolabs) was used to prepare this construct. Primers used in this study are listed in the Key Resource Table.

Selenomethionine-labeled (SeMet) TgMcrB$^{AAA}$ was expressed in minimal medium using methionine auxotrophs (T7 Express Crystal Competent *E. coli*, New England Biolabs) according to manufacturer's protocols. For the expression of native TgMcrB$^{AAA}$, the construct was transformed into *E. coli* BL21(DE3) cells, which were grown at 37°C in Terrific Broth. When OD600 reached 1.0, protein expression was induced by addition of 0.3 mM isopropyl β-D-thiogalactoside (IPTG) and cells were grown overnight at 19°C. Cells were harvested by centrifugation at 6,000x g for 15 minutes at 25°C, and washed twice with Nickel-Loading Buffer (NLB; 20 mM HEPES, pH 7.5, 500 mM NaCl, 30 mM imidazole, 5% glycerol (v/v) and 5 mM β-mercaptoethanol). Pellets were typically flash frozen in liquid nitrogen and stored at -80°C.

Thawed pellets from 500-mL cultures were resuspended in 30 mL of NLB supplemented with 10 mM PMSF, 5 µg/mL DNase I (Roche), 5 mM $MgCl_2$ and a tablet of complete protease inhibitor cocktail (Roche). Lysozyme was added to a final concentration of 1 mg/mL and the mixture was incubated for 15 minutes at 4°C with rocking. Cells were disrupted by sonication and the lysate was cleared of debris by centrifugation at 19,700x g for 30 minutes at 4°C. The supernatant was filtered using a 0.45-µm cut-off syringe filter, incubated at 65°C for 20 minutes, centrifuged at 6,000× g for 15 minutes at 4°C and loaded onto a 5-mL HiTrap chelating column (GE Healthcare) charged with $NiSO_4$ and then washed with NLB. TgMcrB$^{AAA}$ was eluted with an imidazole gradient from 30 mM to 1 M. Peak fractions were pooled, HRV 3C protease was added, and the sample was dialyzed overnight at 4°C against Cleaning Buffer (20 mM HEPES, pH 7.5, 50 mM NaCl, 5% glycerol (v/v) and 5 mM β-mercaptoethanol (10 mM for SeMet-labeled protein)). Another 5-mL HiTrap chelating column charged with $NiSO_4$ was equilibrated with Cleaning Buffer and the sample was applied to this column, followed by elution with a NLB to remove the cleaved 6xHis tag. Pooled peak fractions were concentrated to 2 mL with a centrifugal concentrator (50 kDa cut-off, Millipore). The concentrated protein was further purified by size-exclusion chromatography (SEC) using a HiLoad 16/600 Superdex 200 pg column (GE Healthcare). During

SEC, all proteins were exchanged into $SEC_{150}$ Buffer (20 mM HEPES, pH 7.5, 150 mM KCl, 5 mM $MgCl_2$ and 1 mM DTT (5 mM for SeMet $TgMcrB^{AAA}$)) and then concentrated to 40-80 mg/mL. Concentrations of purified proteins were determined by SDS-PAGE with BSA standards. All point mutations were introduced into $TgMcrB^{AAA}$ in the pET15bp vector by quick-change PCR and the proteins were purified as described above.

**Cloning, expression and purification of TgMcrB**

The gene for full-length TgMcrB (residues 1-613) was amplified by PCR and cloned into the pET15bp vector. The construct was transformed into *E. coli* BL21(DE3) cells, which were grown at 37°C in Terrific Broth. When OD600 reached 1.5, protein expression was induced with 0.3 mM IPTG and cells were grown overnight at 19°C. Cells were harvested and washed twice with NLB. Pellets were typically flash frozen in liquid nitrogen and stored at -80°C. Thawed pellets from 2-L cultures were resuspended in 30 mL of NLB supplemented with 10 mM PMSF, 5 µg/mL DNase I, 5 mM $MgCl_2$ and a tablet of complete protease inhibitor cocktail. Cells were lysed and the full-length TgMcrB protein was purified as described above with the slight modification of using 250 mM KCl in the SEC Buffer (20 mM HEPES, pH 7.5, 250 mM KCl, 5 mM $MgCl_2$ and 1 mM DTT). The protein was concentrated to 20-40 mg/mL.

**Cloning, expression and purification of TgMcrC**

The gene for the *T. gammatolerans EJ3* McrC protein (DOE IMG/M ID 644807739) was codon optimized for expression in *E. coli* and synthesized commercially by Integrated DNA Technologies. The DNA encoding full-length TgMcrC (residues 1-458) was amplified by PCR and cloned into the pCAV6 vector, a modified version of the pMAL c5x T7 expression vector, in which the Factor Xa cleavage site upstream of the TgMcrC gene was replaced with an HRV 3C cleavage site and a 6xHis tag was introduced upstream of the MBP sequence. Gibson assembly was used to prepare this construct.

The TgMcrC construct was transformed into *E. coli* BL21(DE3) cells, which were grown at 37°C in Terrific Broth. When OD600 reached 1.0, protein expression was induced with 0.3 mM IPTG and the cells were grown overnight at 19°C. Cells were harvested and washed twice with NLB. Pellets were typically flash frozen in liquid nitrogen and stored at -80°C. Thawed pellets from 500-mL cultures were resuspended in 30 mL of NLB supplemented with 10 mM PMSF, 5 µg/mL DNase I, 5 mM $MgCl_2$ and a tablet of complete protease inhibitor cocktail. Lysozyme was added to a final concentration of 1 mg/mL and the mixture was incubated for 15 minutes at 4°C with rocking. Cells were disrupted by sonication and the lysate was cleared of debris by centrifugation at 19,700x g for 30 minutes at 4°C. The supernatant was filtered using a 0.45-μm cut-off syringe filter, loaded onto a 5-mL HiTrap chelating column charged with $NiSO_4$ and then washed with NLB. TgMcrC was eluted with an imidazole gradient from 30 mM to 1 M. Peak fractions were pooled, HRV 3C protease was added, and the sample was dialyzed overnight at 4°C against SP-Loading Buffer (SPLB; 20 mM HEPES, pH 7.5, 250 mM NaCl, 1 mM EDTA, 5% glycerol (v/v) and 1 mM DTT). The sample was applied to a 5-mL HiTrap SP HP column (GE Healthcare) equilibrated with SPLB and then washed with SPLB. TgMcrC was eluted with a NaCl gradient from 250 mM to 1 M. Because TgMcrC is prone to precipitate, no further purification steps were attempted and the pooled peak fractions yielded protein at a purity of ~70% and a concentration of ~0.8 mg/mL. All point mutations were introduced into TgMcrC in the pCAV6 vector by quick-change PCR and the proteins were purified as described above.

**Cloning, expression and purification of EcMcrB**

The gene for full-length *E. coli* McrB (Uniprot P15005; DOE IMG ID 646316336) was codon optimized for expression in *E. coli* and synthesized commercially by GENEART. The DNA encoding full-length EcMcrB (residues 1-459) was cloned into the pMAL-c2Xp vector, a modified version of the pMAL-c2X vector (New England Biolabs), in which the Factor Xa cleavage site after the N-terminal MBP tag was replaced with an HRV 3C cleavage site.

EcMcrB was transformed into *E. coli* BL21(DE3) cells, which were grown at 37°C in Terrific Broth. When OD600 reached 1.0, protein expression was induced with 0.3 mM IPTG and cells were grown overnight at 19°C. Cells were harvested and washed once with $TGED_{500}$ Buffer (20 mM Tris-HCl, pH 8.0, 5% glycerol (v/v), 1 mM EDTA, 1 mM DTT and 500 mM NaCl). Pellets were flash frozen in liquid nitrogen and stored at -80°C. Thawed pellets from 500-mL cultures were resuspended in 30 mL of $TGED_{500}$ Buffer supplemented with 10 mM PMSF, 5 μg/mL DNase I, 5 mM $MgCl_2$, and a tablet of complete protease inhibitor cocktail. Lysozyme was added to a final concentration of 1 mg/mL and the mixture was incubated for 15 minutes at 4°C with rocking. Cells were disrupted by sonication and the lysate was cleared of debris by centrifugation at 19,700× g for 30 minutes at 4°C. The supernatant was filtered using a 0.45-μm cut-off syringe filter, loaded onto a 5-mL HiTrap MBP column (GE Healthcare), washed with $TGED_{500}$ and eluted with 10 mM D-maltose in $TGED_{500}$ Buffer. Peak fractions were pooled, HRV 3C protease was added, and the sample was dialyzed overnight at 4°C against $TGED_{50}$ Buffer ($TGED_{500}$ Buffer but with 50 mM NaCl instead of 500 mM). The sample was then applied to a 5-mL HiTrap Q HP ion-exchange column (GE Healthcare) in $TGED_{50}$ and eluted with a NaCl gradient from 50 mM to 500 mM. Peak fractions were pooled, concentrated and further purified by SEC using a HiLoad 16/600 Superdex

200 pg column, during which the protein was exchanged into $SEC_{150}$ Buffer. The protein was then concentrated to ~25 mg/mL.

**Cloning, expression and purification of EcMcrC**

The gene encoding full-length *E. coli* McrC protein (Uniprot P15006; DOE IMG ID 637004274) was codon optimized for expression in *E. coli* and synthesized commercially by GENEART. The DNA encoding full-length EcMcrC (residues 1-348) was cloned into the pMAL-c2Xp vector.

EcMcrC was transformed into *E. coli* BL21(DE3) cells and grown at 37°C in Terrific Broth. When OD600 reached 1.0, protein expression was induced with 0.3 mM IPTG and cells were grown overnight at 19°C. Cells were harvested and washed once with $TGED_{500}$ Buffer. Pellets were flash frozen in liquid nitrogen and stored at -80°C. Thawed pellets from 500-mL cultures were resuspended in 30 mL of $TGED_{500}$ Buffer supplemented with 10 mM PMSF, 5 µg/mL DNase I, 5 mM $MgCl_2$, and a tablet of complete protease inhibitor cocktail. Lysozyme was added to a final concentration of 1 mg/mL and the mixture was incubated for 15 minutes at 4ºC with rocking. Cells were disrupted by sonication and the lysate was cleared of debris by centrifugation at 19,700x g for 30 minutes at 4ºC. The supernatant was filtered using a 0.45-µm cut-off syringe filter, loaded onto a 5-mL HiTrap MBP column, washed with $TGED_{500}$ Buffer and eluted with 10 mM D-maltose in $TGED_{500}$ Buffer. Peak fractions were pooled, HRV 3C protease was added, and the sample was dialyzed overnight at 4ºC against $HGED_{250}$ Buffer (20 mM HEPES, pH 7.5, 5% glycerol (v/v), 1 mM EDTA, 1 mM DTT and 250 mM NaCl). The sample was then applied to a 5-mL HiTrap SP HP ion-exchange column in $TGED_{250}$ Buffer and eluted with a NaCl gradient from 250 mM to 1 M. Because EcMcrC is prone to precipitate, no further purification steps were attempted. The pooled peak fractions yielded protein at a purity of ~70% and a concentration of ~6 mg/mL.

**GTPase activity assays**

GTPase activity was measured by using a colorimetric Malachite green assay that monitors the amount of free phosphate released over time (Leonard et al., 2005). To measure the basal GTPase activity of $TgMcrB^{AAA}$, 0.4 µM $TgMcrB^{AAA}$ was incubated with 1 mM GTP at 65ºC in Reaction Buffer (20 mM Tris-HCl, pH 8.0, 150 mM KCl 5 mM $MgCl_2$). To measure the GTPase activity of $TgMcrB^{AAA}$ stimulated by TgMcrC, the same conditions were used but 0.1 µM TgMcrC was added. At time points of 0, 5, 10, 20, 30, 45, 60, 80, 100 and 120 minutes, 20-µL aliquots were taken and quenched with 5 µL of 0.5 M ETDA, pH 8.0. For colorimetric reactions, 150 µL of filtered Malachite green solution were added to each sample and incubated for 5 minutes. The absorbance at 650 nm of the samples was measured with a Multiskan GO Microplate Spectrophotometer (Thermo Scientific). The amount of phosphate released was determined using a standard curve. The specific activity is reported for all wildtype and mutant proteins. Quantified data represent the average of three independent experiments using multiple independently purified batches of protein with error bars indicating the standard deviation from the mean (n=3, mean ± standard deviation).

**Negative-stain EM**

Negatively stained samples were prepared as described (Ohi et al., 2004). Freshly purified proteins were diluted to ~0.05 mg/mL with $SEC_{150}$ Buffer supplemented with 2.5 mM GTPγS before applying 5-µL aliquots to glow-discharged grids. Grids were stained with 0.7% uranyl formate (Pfaltz & Bauer, U01000) and imaged with a Philips CM10 electron microscope equipped with a tungsten filament and operating at 100 kV. All images were recorded on an AMT XR16L-ActiveVu charge-coupled device camera (Woburn, MA, USA) using a defocus of approximately -1.5 µm and a nominal magnification of 52,000x.

**Crystallization, X-ray data collection and structure determination of $TgMcrB^{AAA}$**

SeMet $TgMcrB^{AAA}$ with 2.5 mM GTPγS was crystallized by sitting drop vapor diffusion in 0.1 M sodium acetate, pH 6.5, 17.5% 2-methyl-2,4-pentanediol (v/v) with a drop size of 2 µL and a reservoir volume of 650 µL. Crystals appeared within 3-4 days at 20°C and were cryo-protected with Parabar 10312 (Hampton Research) and frozen in liquid nitrogen. Single-wavelength anomalous diffraction (SAD) data were collected remotely on the tunable NE-CAT 24-ID-C beamline at the Advanced Photon Source at the selenium edge energy of 12.663 keV (0.9791 Å) (Supplementary Table S1). Data were integrated and scaled using the NE-CAT RAPD pipeline. Strong anomalous signal was obtained from a single crystal diffracting to 2.9 Å (space group P2; unit cell dimensions: a = 100.24 Å, b = 108.87 Å, c = 118.67 Å and α = 90.00º, β = 107.41º, γ = 90.00º) and multiple SeMet SAD datasets were collected from different positions of this crystal. All possible combinations of datasets were tested and merged using the program BLEND in the CCP4 suite (Foadi et al., 2013; Winn et al., 2011). Experimental phases were obtained from the combination with the strongest anomalous signal. Heavy-atom sites were located using SHELX C/D/E (Sheldrick, 2010) and phasing, density modification, and initial model building were carried out using the CRANK2 pipeline (Skubák and Pannu, 2013). Iterative rounds of refinement and model building were carried out using the programs COOT (Emsley et al., 2010) and REFMAC (Murshudov et al., 2011) to improve the initial model, which resolved most regions of four

$TgMcrB^{AAA}$ monomers. This partial model was used as the search model to perform molecular replacement using PHASER (Bunkóczi et al., 2013) on a native dataset diffracting to 2.83 Å (space group P2; unit cell dimensions: a = 100.02 Å, b = 108.55 Å, c = 118.43 Å and α = 90.00º, β = 106.94º, γ = 90.00º), which was collected at the NE-CAT 24-ID-E beamline at the selenium edge energy of 12.663 keV (0.9791 Å). Further model building and refinement was carried out manually in COOT and PHENIX, respectively (Adams et al., 2010; Emsley et al., 2010). The final model contained four well-resolved and two poorly-resolved molecules in the asymmetric unit and was refined to 2.95 Å resolution with $R_{work}/R_{free}$ values of 0.345 / 0.364 (Supplementary Table S1).

**Cryo-EM sample preparation and data collection**

For $TgMcrB^{AAA}$, thawed protein was diluted to 10 mg/mL with $SEC_{150}$ Buffer containing 2.5 mM GTPγS. Samples were mixed with 20x digitonin (Calbiochem) stock to a final concentration of 0.05%, and 3.5 µL aliquots were applied to C-flat thick holey carbon grids (CF-1.2/1.3-4C-T, Protochips), blotted for 7 seconds at 4°C and plunge-frozen in liquid ethane using a Vitrobot Mark IV (Thermo Fisher Scientific).

For the $TgMcrB^{AAA}C$ complex, thawed $TgMcrB^{AAA}$ was mixed with freshly purified TgMcrC at a molar ratio of 4:1. The sample was concentrated using a 2-mL centrifugal concentrator (100 kDa cut-off, Millipore). Concentrated protein was buffer-exchanged to $SEC_{150}$ Buffer in the concentrator and the final concentration was estimated to be ~14 mg/mL. The complex was then mixed with 50x GTPγS stock solution to a final GTPγS concentration of 2.5 mM and incubated for 30 minutes at 4ºC. Samples were mixed with 20x digitonin to a final concentration of 0.05%, and 3.5-µL aliquots were applied to C-flat thick holey carbon grids (CF-1.2/1.3-4C-T), blotted for 8-10 seconds at 4ºC and plunge-frozen in liquid ethane using a Vitrobot Mark IV.

For the TgMcrBC and EcMcrBC complexes, thawed McrB was mixed with freshly purified McrC at a molar ratio of 4:1. The samples were concentrated using 2-mL centrifugal concentrators (100 kDa cut-off, Millipore). Concentrated proteins were then buffer-exchanged into $SEC_{250}$ Buffer (for TgMcrBC) or $SEC_{150}$ Buffer (for EcMcrBC) in the concentrators and finally concentrated to ~16 mg/mL. The prepared complexes were mixed with 50x GTPγS stock solution to a final GTPγS concentration of 2.5 mM and incubated for 30 minutes at 4ºC. Samples were mixed with 20x digitonin to a final concentration of 0.05% digitonin, and 3.5 µL aliquots were applied to Quantifoil R1.2/1.3 400 mesh Au grids, blotted for 8-10 seconds at 4ºC and plunge-frozen in liquid ethane using a Vitrobot Mark IV.

Cryo-EM data were collected on a 300-kV Titan Krios electron microscope (Thermo Fisher Scientific) equipped with a K2 Summit direct electron detector at a nominal magnification of

29,000x in super-resolution counting mode. After binning over 2 x 2 pixels, the calibrated pixel size was 1.0 Å on the specimen level. For all specimens other than EcMcrBC, exposures of 10 seconds were dose-fractionated into 40 frames with a dose rate of 8 electrons per pixel per second, resulting in a total dose of 80 electrons per Å$^2$. For EcMcrBC, exposures of 20 seconds were dose-fractionated into 40 frames with a dose rate of 4 electrons per pixel per second, resulting in a total dose of 80 electrons per Å$^2$. Cryo-EM data collection statistics are summarized in Supplementary Table S2.

**Cryo-EM data processing**

For TgMcrB$^{AAA}$ and the TgMcrB$^{AAA}$C complex, image processing was done in RELION-3.0-beta (Nakane et al., 2018; Zivanov et al., 2018; Zivanov et al., 2019) and images for TgMcrBC and EcMcrBC were processed in both CryoSPARC-2.4.0 (Structura Biotechnology) (Punjani et al., 2017) and RELION-3. All movie frames were corrected with a gain reference collected during the same EM session, and specimen movement was corrected using RELION's implementation of motion correction (for TgMcrB$^{AAA}$ and TgMcrB$^{AAA}$C) or MotionCorr2 (for TgMcrBC and EcMcrBC) with dose weighting (Zheng et al., 2017; Zivanov et al., 2018). The contrast transfer function (CTF) parameters were estimated using CTFFIND-4 (Rohou and Grigorieff, 2015) for TgMcrB$^{AAA}$ and TgMcrB$^{AAA}$C or Gctf-1.0.6 (Zhang, 2016) for TgMcrBC and EcMcrBC. Images showing substantial ice contamination, abnormal background, thick ice, low contrast or poor Thon rings were discarded.

For TgMcrB$^{AAA}$, 1,599 micrographs were collected, of which 1,517 micrographs were selected for further processing. Particles were picked with Gautomatch (https://www.mrc-lmb.cam.ac.uk/kzhang/Gautomatch/) without templates, which identified 277,503 particles that were windowed into 320x320-pixel images. The particle images were binned 4 times and subjected to two rounds of 2D classification. Classes that produced averages with fine structural detail and no overlap with neighboring particles were combined and used to generate an initial reference map. The selected 153,891 particles were subjected to 3D classification into four classes, three of which were selected and used to re-extract the corresponding particles into 320x320-pixel images that were then rescaled into 256x256-pixel images. The centered, re-extracted particles were refined with C1 symmetry to a resolution of 3.4 Å, according to the Fourier shell correlation (FSC) = 0.143 criterion (Rosenthal and Henderson, 2003), which was used for all resolution estimates. Subsequent CTF refinement and Bayesian polishing improved the overall resolution of the map to 3.1 Å.

For the TgMcrB$^{AAA}$C complex, 1,795 of the 2,070 collected micrographs were selected for further processing. Gautomatch was used to pick the first 200 micrographs without templates, and the ~10,000 picked particles were subjected to 2D classification. Four representative class averages were then selected as templates for Gautomatch to pick particles from all the micrographs. The 264,850 auto-picked particles were cleaned up by two rounds of 2D classification. The particles from eight classes with well-defined averages (156,149 particles) were used to generate an initial density map in RELION, which was then used as reference for 3D classification of the cleaned-up particles into six classes. Four classes showed good fine structure and were combined (115,774 particles), and subsequent 3D refinement with C1 symmetry, CTF refinement and Bayesian polishing yielded a map at 4.4-Å resolution. A second dataset collected using the same conditions was processed following the same strategy, yielding a map at 4.3-Å resolution from 88,819 refined particles. The particles from the two datasets were combined and further refined with C1 symmetry to generate an improved map at 4.2 Å (204,593 particles). While this map showed strong density for one half of the complex, the other half was represented by substantially weaker density. 3D refinement was thus repeated with C2 symmetry imposed, which yielded a symmetrized map for the full complex map at 4.2-Å resolution. To overcome the flexibility of the connection between the two half-complexes, particles in the non-symmetrized map were subjected to automated multi-body refinement implemented in RELION-3, using individual masks for the two half-complexes that overlapped in the region of the two-fold axis. Signal subtraction was performed for each rigid body using the 'relion_flex_analyse' command, which only retains the signal inside the selected rigid body (Nakane et al., 2018). The signal-subtracted particles for one of the two bodies were used to calculate a reference map for the half-complex using the 'relion_reconstruct' command. The signal-subtracted particles for both bodies were combined (409,186 particles) and subjected to 3D refinement with C1 symmetry and starting with a global search. Subsequent CTF refinement and 3D refinement yielded the final map for the half-complex at an overall resolution of 3.7 Å.

For the TgMcrBC complex, 1,936 of 2,078 micrographs and for the EcMcrBC complex, 1,088 of 1,161 micrographs were selected for further processing. Particles were picked with Gautomatch with templates generated from preliminary data collected on a 200-kV Talos Arctica electron microscope (Thermo Fisher Scientific). The auto-picked particles (354,707 for the TgMcrBC complex and 184,487 for the EcMcrBC complex) were extracted into 320x320-pixel images that were then rescaled into 256x256-pixel images. All particle images were used for *ab-initio* reconstruction in Cryosparc-2.4.0, specifying three output classes. The best of the three maps, including 226,813 particles for the TgMcrBC complex and 106,684 particles for the

EcMcrBC complex, were selected for non-uniform refinement, which yielded maps for the half-complexes at 3.0-Å resolution for the TgMcrBC complex and at 4.9-Å resolution for the EcMcrBC complex. The particles were transferred back to RELION using the pyem package (https://github.com/asarnow/pyem), re-extracted into 320x320-pixel images and further refined without imposing symmetry to generate maps for the TgMcrBC complex at 2.9-Å resolution and for the EcMcrBC complex at 4.1-Å resolution. CTF refinement and Bayesian polishing improved the maps to resolutions of 2.7 and 3.5 Å, respectively. Finally, the particles were re-extracted into 400x400-pixel images. Refinement, CTF refinement and Bayesian polishing yielded the final maps at 2.4-Å resolution for TgMcrBC and at 3.3-Å resolution for EcMcrBC.

**Model building and refinement**

For the TgMcrB$^{AAA}$ hexamer, the best refined monomer from the X-ray model was fit into each subunit density of the 3.1-Å resolution cryo-EM map using UCSF chimera (Pettersen et al., 2004). Further iterative refinement cycles between the phenix.real_space_refine command in PHENIX with secondary structure restraints and manual adjustments in COOT yielded the final model for the TgMcrB$^{AAA}$ hexamer.

For the TgMcrB$^{AAA}$C complex, the final cryo-EM model of the TgMcrB$^{AAA}$ hexamer was manually fit into the density map of the half-complex and refined using the phenix.real_space_refine command in PHENIX with morphing, simulated annealing and secondary structure restraints. *Ab-initio* model building for TgMcrC was carried out in COOT (Emsley et al., 2010), guided by secondary structure predictions from SPIDER2 (Yang et al., 2017) and PSIPRED (Buchan et al., 2013). The density for TgMcrC was good up to residue 312, but the remaining C-terminal endonuclease domain was poorly resolved. Therefore, a homology search was performed in I-TESSAR (Roy et al., 2010; Yang et al., 2015; Yang and Zhang, 2015) for TgMcrC residues 312-458, and a homology model was generated based on the *Saccharolobus solfataricus* holiday junction resolving enzyme (PDB: 1OB8; Middleton et al., 2004). This homology model was fit into the corresponding density and manual adjustments were performed in COOT. Finally, all built models were combined and iterative cycles of real-space refinement in PHENIX with secondary structure restraints and manual adjustments in COOT were performed, yielding the final model for the TgMcrB$^{AAA}$C half-complex.

For the TgMcrBC complex, the final cryo-EM model of TgMcrB$^{AAA}$C was manually fit into the density map of the half-complex and refined using the phenix.real_space_refine command in PHENIX with morphing, simulated annealing and secondary structure restraints. Because the nuclease domain of TgMcrC was poorly resolved, most regions were removed from the model.

Finally, iterative cycles of real-space refinement in PHENIX with secondary structure restraints and manual adjustments in COOT were performed, yielding the final model of the TgMcrBC half-complex.

For the EcMcrBC complex, SWISS-MODEL (Waterhouse et al., 2018) was used to generate a homology model of EcMcrB based on the $TgMcrB^{AAA}$ structure, which was manually fit into one subunit in the 3.3-Å cryo-EM map using UCSF chimera. After manual adjustments in COOT, the corrected model was fit into each subunit of the hexameric EcMcrB density. To build the EcMcrC model, the unique N-terminal domain was removed from the TgMcrC model, and all the residues were mutated to alanine except for the highly conserved residues (based on the sequence alignment of McrC homologs). This model was manually fit into the corresponding density of the 3.3-Å resolution cryo-EM map in UCSF chimera, followed by manual adjustment of each residue in COOT. Manual adjustment was guided by secondary structure predictions from SPIDER2 and PSIPRED. Finally, all built models were combined, and iterative cycles of real-space refinement in PHENIX with secondary structure restraints and manual adjustments in COOT yielded the final model of the EcMcrBC half-complex.

All the refinement statistics are summarized in Supplementary Table S2. For model validation, the final model for each map was refined against one of the independent half maps (map 1) of the corresponding map. FSC curves were then calculated between the refined model and half map 1 (work), half map 2 (free) as well as the combined map (Figure S1h, S2f, S4f and S6g).

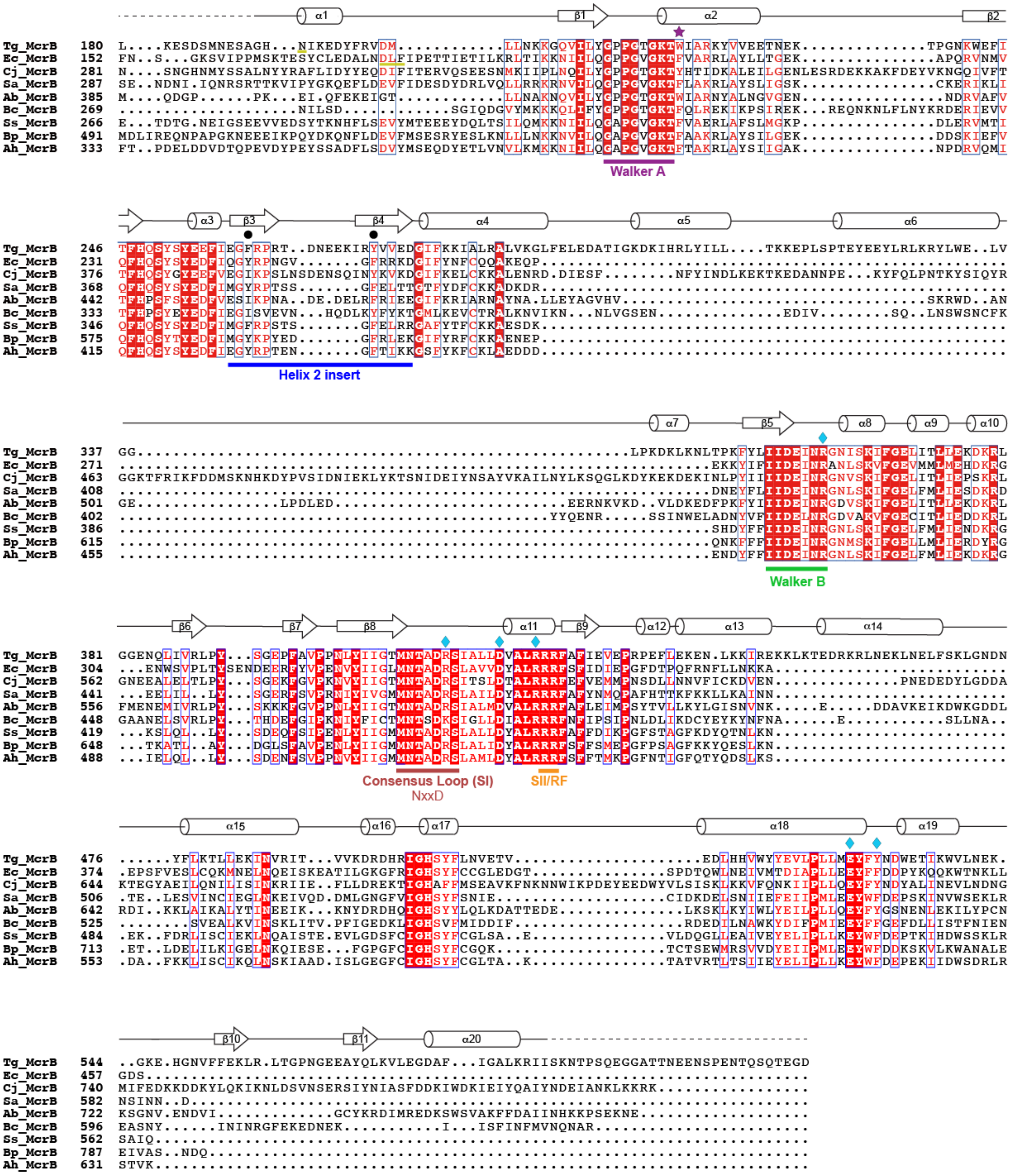


### Supplementary Data S1

**Supplementary Data S1. Sequence alignment of McrB family proteins.** Conserved motifs are labeled and colored. The secondary structure diagram is based on the AAA domain of TgMcrB. Symbols denote the following: cyan diamonds, interface residues contributing to tight/loose asymmetry; black circles, hydrophobic residues that stabilize the loop-helix-loop region of the McrC finger domain; purple star, π-stacking residue that contacts the guanine base. Segments near α1 involved in guanine nucleotide recognition are underlined in gold in Tg and EcMcrB sequences. Abbreviations are as follows: Tg, *Thermococcus gammatolerans*; Ec, *Escherichia coli*; Cj, *Campylobacter jejuni*; Sa, *Staphylococcus aureus*; Ab, *Aciduliprofundum boonei*; Bc, *Bacillus cereus*; Ss, *Streptococcus suis*; Bp, *Butyrivibrio proteoclasticus*; Ah, *Anaerobutyricum hallii*.

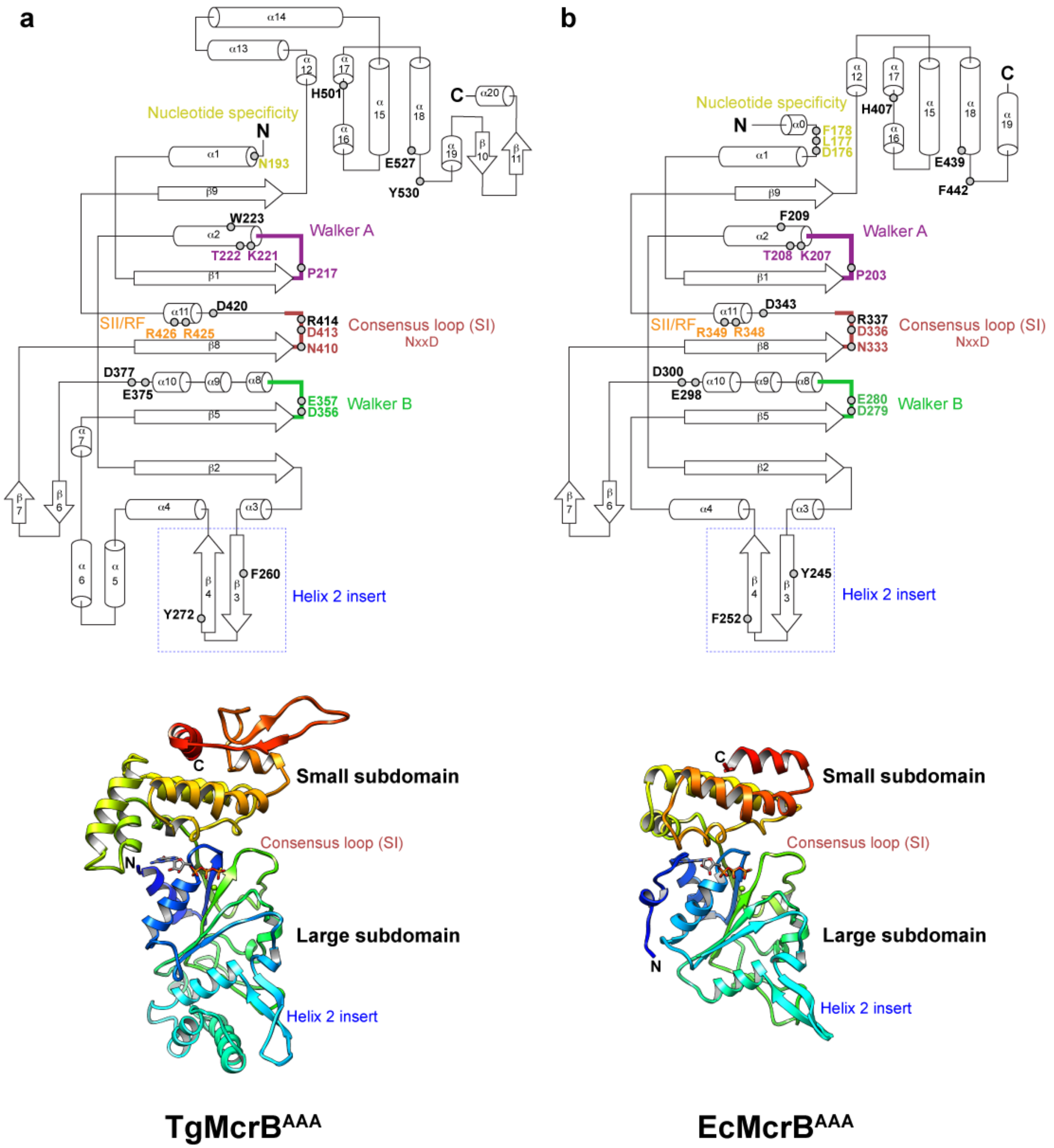


### Supplementary Data S2

**Supplementary Data S2. Secondary structure topology diagram and structures of the monomeric AAA+ domains of TgMcrB (a) and EcMcrB (b) in ribbon representation.** Conserved motifs among McrB family proteins are labeled and colored as in Supplementary Data S1.

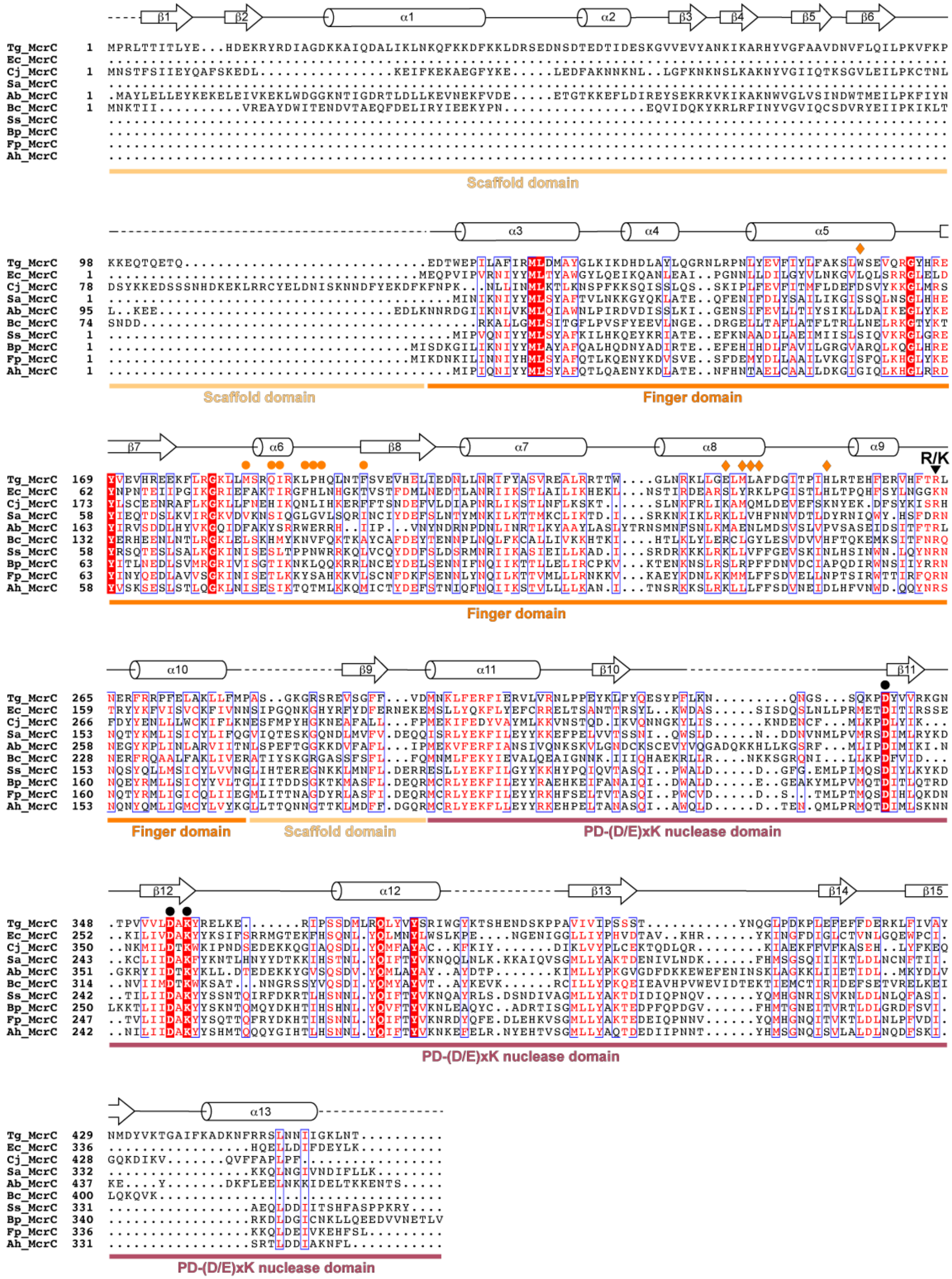


**Supplementary Data S3**

**Supplementary Data S3. Sequence alignment of McrC family proteins.** Symbols denote the following: orange circle, loop-helix-loop residues interacting with hydrophobic side chains at the base of the McrB ring (Figure 3e); orange diamonds, residues that form anchoring interactions at the distorted tight McrB interface (Figure 3f); black triangle, conserved arginine or lysine whose side chain is important for the stimulation of McrB GTPase activity; black circles, residues critical for McrC nuclease activity. The secondary structure diagram is based on TgMcrC. Individual domain segments are colored as in Figure 3d and labeled below the alignment. Abbreviations are the same as in Supplementary Data S1.

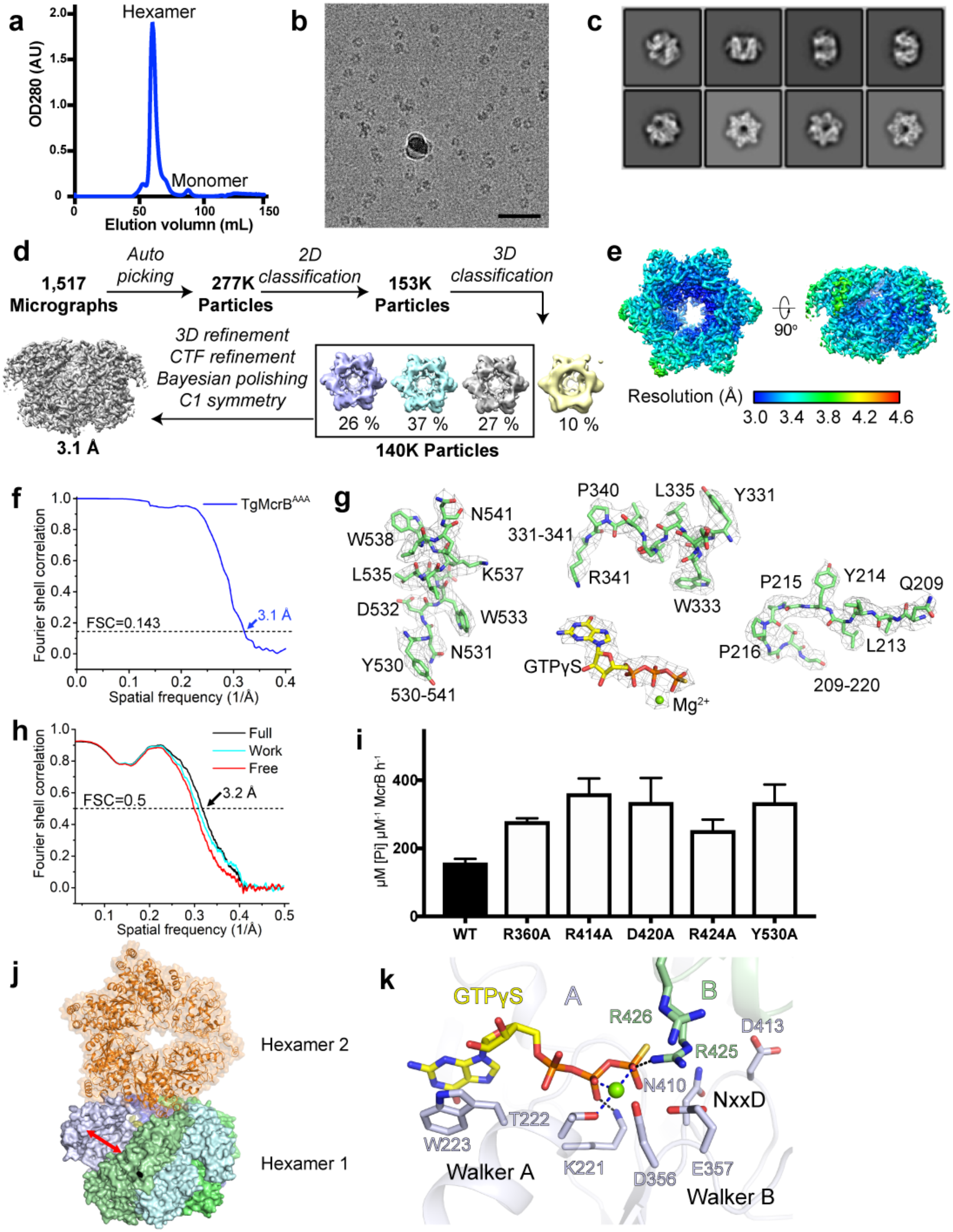
a
Hexamer
Monomer
OD280 (AU)
Elution volumn (mL)
b
c
d
1,517 Micrographs
Auto picking
277K Particles
2D classification
153K Particles
3D classification
3D refinement
CTF refinement
Bayesian polishing
C1 symmetry
26 %
37 %
27 %
10 %
140K Particles
3.1 Å
e
90°
Resolution (Å)
3.0 3.4 3.8 4.2 4.6
f
Fourier shell correlation
Spatial frequency (1/Å)
FSC=0.143
3.1 Å
g
N541
W538
331-341
K537
L535
D532
W533
N531
Y530
530-541
P340
L335
Y331
R341
W333
GTPγS
Mg2+
P215
Y214
Q209
P216
L213
209-220
h
Full
Work
Free
FSC=0.5
3.2 Å
i
μM [Pi] μM-1 McrB h-1
WT
R360A
R414A
D420A
R424A
Y530A
j
Hexamer 2
Hexamer 1
k
GTPγS
A
B
R426
D413
R425
N410
NxxD
T222
W223
Walker A
K221
D356
E357
Walker B

1

**Figure S1**

**Figure S1: Biochemical and structural analysis of the TgMcrB$^{AAA}$ hexamer, related to Figures 1 and 2.** (**a**) Gel-filtration profile of the TgMcrB$^{AAA}$ hexamer. (**b**) Cryo-EM image area of vitrified TgMcrB$^{AAA}$ hexamer. Scale bar is 50 nm. (**c**) Selected 2D-class averages of the TgMcrB$^{AAA}$ hexamer. Side length of the individual panels is 32 nm. (**d**) Cryo-EM data-processing workflow for the TgMcrB$^{AAA}$ hexamer. (**e**) Local resolution map for the TgMcrB$^{AAA}$ hexamer. (**f**) Gold-standard Fourier shell correlation (FSC) curve for the TgMcrB$^{AAA}$ hexamer after correction for masking effects. The resolution was estimated based on the FSC = 0.143 criterion. (**g**) Cryo-EM densities for selected regions in the map of the TgMcrB$^{AAA}$ hexamer. (**h**) Cross-validation FSC curves for the TgMcrB$^{AAA}$ hexamer: cyan curve, refined model *versus* half map 1 used for refinement (Work); red curve, refined model *versus* half map 2 not used for refinement (Free); black curve, refined model *versus* the combined final map (Full). The similarity of the 'work' and 'free' curves suggests no substantial over-fitting. The correlation is above 0.5 up to a resolution of 3.2 Å. (**i**) GTPase activity of wild-type TgMcrB$^{AAA}$ and mutants in which alanine substitutions were introduced at residues at the tight and loose interfaces shown in Figure 1d and e (n = 3, mean ± standard deviation). (**j**) Structure of the 'open-ring' assembly of the TgMcrB$^{AAA}$ hexamer determined by X-ray crystallography. The subunits in hexamer 1 are colored as in Figure 1, and symmetry-related hexamer 2 is colored in orange. The red double arrow indicates the subunit separation at the A/B interface likely introduced by the crystal packing. (**k**) Close-up view of the GTP-binding site at the loose A/B interface in the cryo-EM structure of the TgMcrB$^{AAA}$ hexamer. Dashed lines indicate hydrogen bonds (black) and metal coordination (blue).

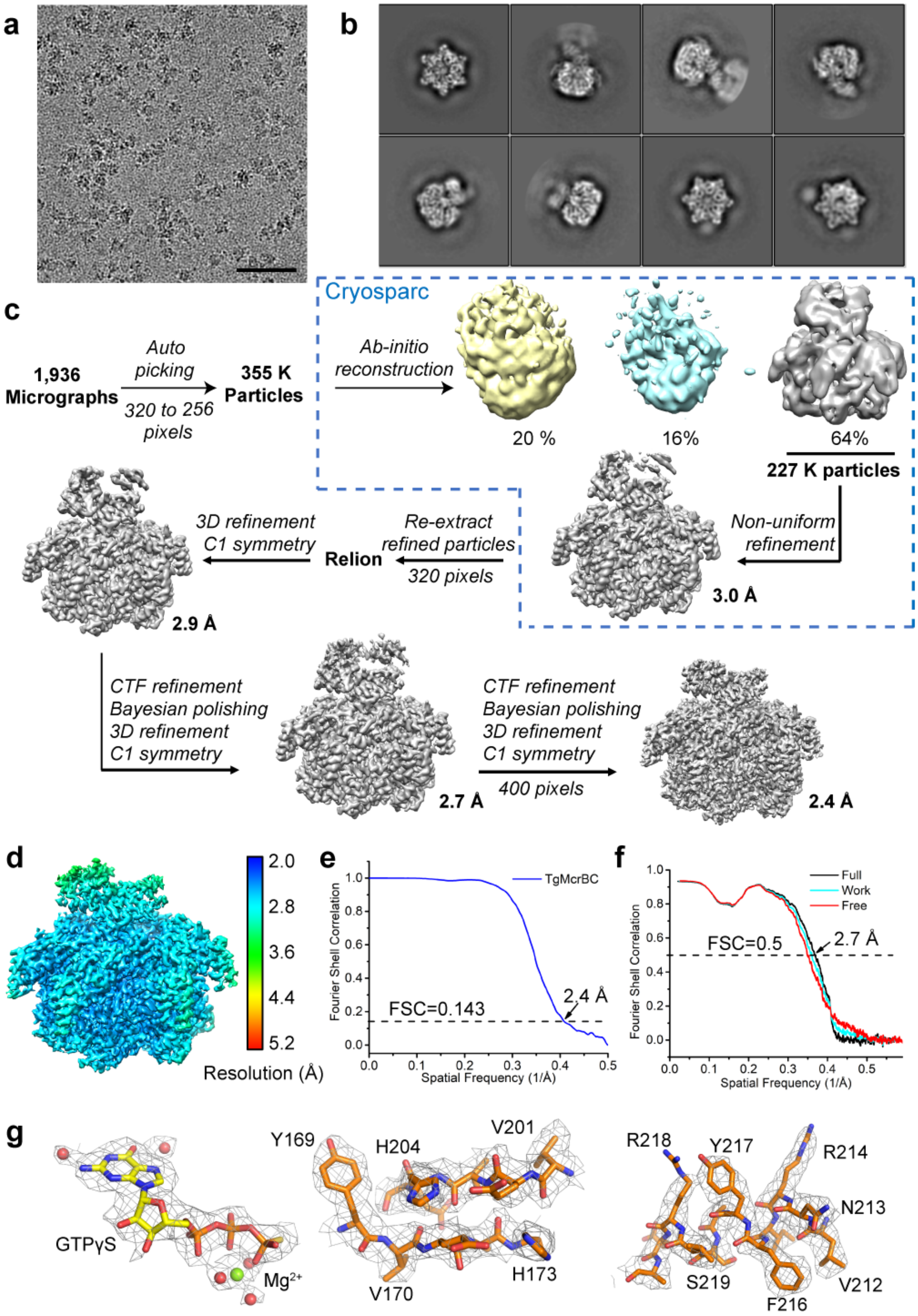

a
b
c
Cryosparc
1,936 Micrographs
Auto picking
320 to 256 pixels
355 K Particles
Ab-initio reconstruction
20 %
16%
64%
227 K particles
Non-uniform refinement
3.0 Å
Re-extract refined particles
320 pixels
Relion
3D refinement C1 symmetry
2.9 Å
CTF refinement
Bayesian polishing
3D refinement
C1 symmetry
2.7 Å
CTF refinement
Bayesian polishing
3D refinement
C1 symmetry
400 pixels
2.4 Å
d
2.0
2.8
3.6
4.4
5.2
Resolution (Å)
e
TgMcrBC
Fourier Shell Correlation
FSC=0.143
2.4 Å
Spatial Frequency (1/Å)
f
Full
Work
Free
FSC=0.5
2.7 Å
Fourier Shell Correlation
Spatial Frequency (1/Å)
g
GTPγS
Mg2+
Y169
H204
V201
V170
H173
R218
Y217
R214
N213
S219
F216
V212


**Figure S2**

1

**Figure S2: Cryo-EM analysis of the TgMcrBC complex, related to Figures 3 and 4.** (**a**) Cryo-EM image area of vitrified TgMcrBC complex. Scale bar is 50 nm. (**b**) Selected 2D-class averages of the TgMcrBC complex. Side length of the individual panels is 32 nm. (**c**) Cryo-EM data-processing workflow for the TgMcrBC complex. (**d**) Local resolution map for the TgMcrBC half-complex. (**e**) Gold-standard FSC curve for the TgMcrBC 'half'-complex after correction for masking effects. The resolution was estimated based on the FSC = 0.143 criterion. (**f**) Cross-validation FSC curves for the TgMcrBC half-complex: cyan curve, refined model *versus* half map 1 used for refinement (Work); red curve, refined model *versus* half map 2 not used for refinement (Free); black curve, refined model *versus* the combined final map (Full). The similarity of the 'work' and 'free' curves suggests no substantial over-fitting. The correlation is above 0.5 up to a resolution of 2.7 Å. (**g**) Cryo-EM densities for selected regions in the map of the TgMcrBC complex.

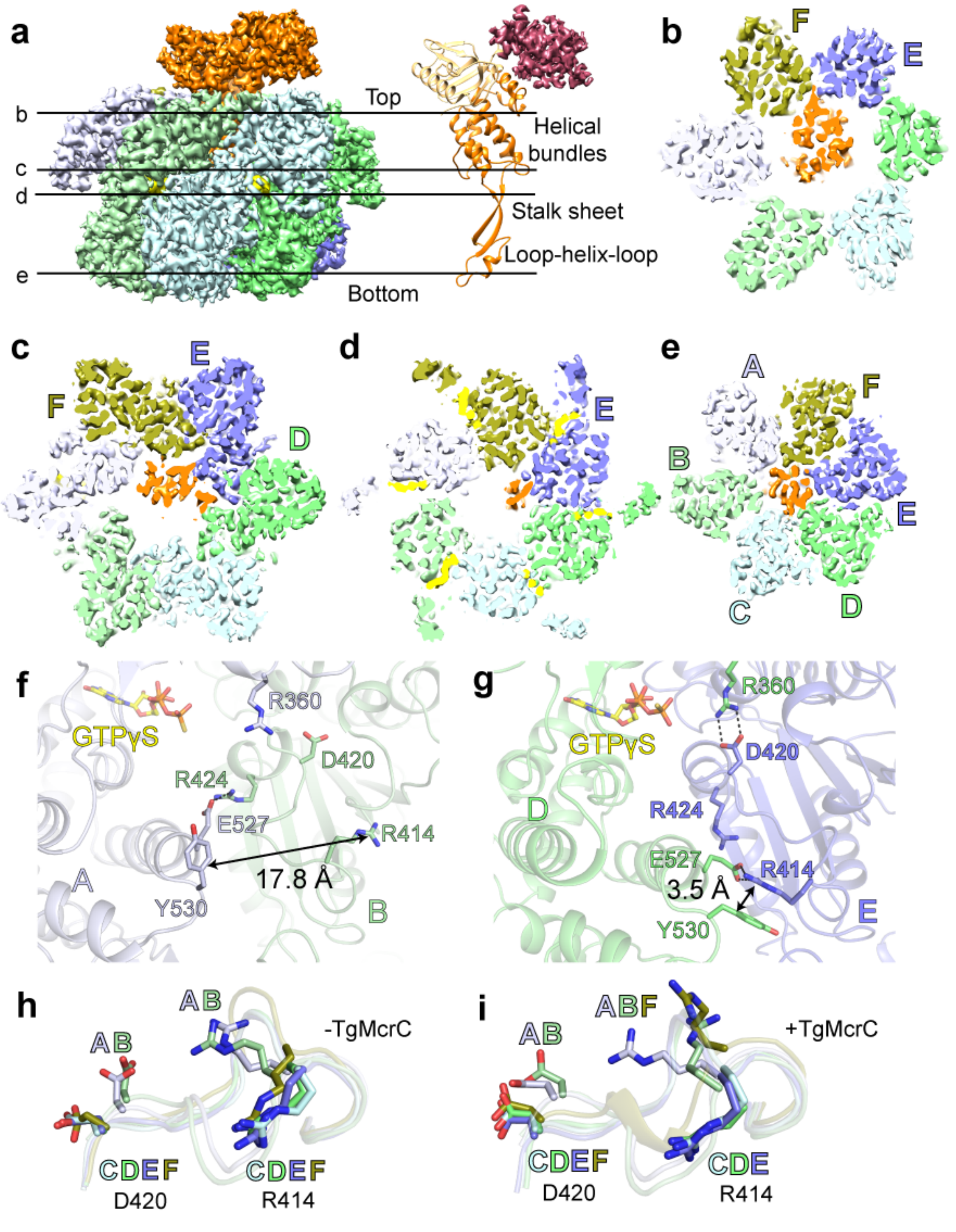


**Figure S3**

**Figure S3: Bound TgMcrC adopts to the intrinsic asymmetry of the TgMcrB hexamer, related to Figure 3 and 4.** (**a**) Side views of the TgMcrBC half-complex in surface representation (left panel) and TgMcrC in cartoon representation (right panel). (**b–e**) Slice sections through the density at the levels indicated by the solid lines in (**a**). The TgMcrB subunits forming the main interactions with TgMcrC at the different levels are labeled. (**f** and **g**) Close-up views of interacting residues at the tight A/B interface (**f**) and the loose D/E interface (**g**) of the TgMcrB hexamer, shown from the same angles as in Figure 1d and e. (**h** and **i**) Superpositions of the 414-420 loop of the six subunits in TgMcrB by itself (**h**) and in TgMcrB in complex with TgMcrC (**i**).

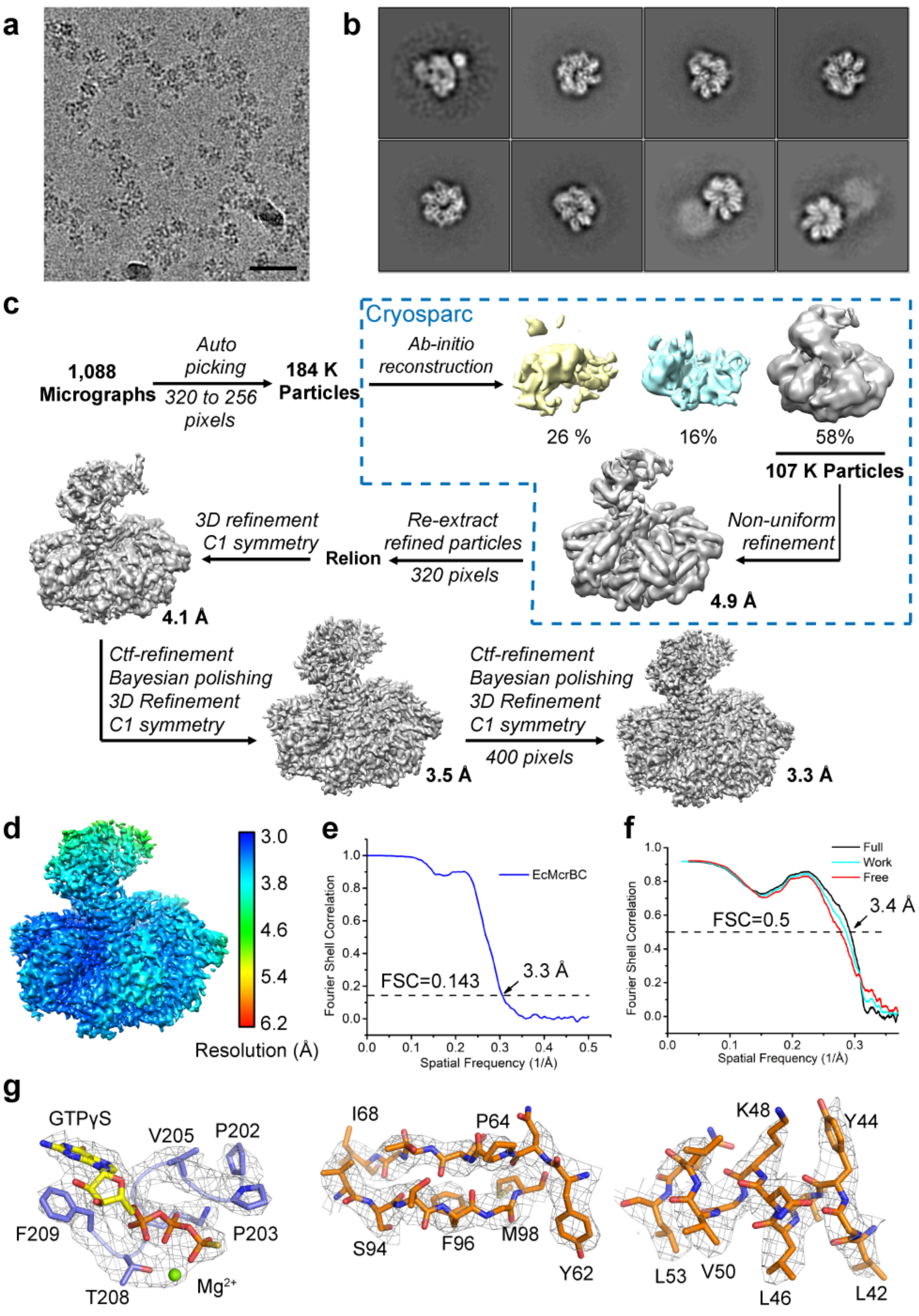


**Figure S4**

**Figure S4: Cryo-EM analysis of the EcMcrBC complex, related to Figure 5**. (**a**) Cryo-EM image area of vitrified EcMcrBC complex. Scale bar is 50 nm. (**b**) Selected 2D-class averages of the EcMcrBC complex. Side length of the individual panels is 32 nm. (**c**) Cryo-EM data-processing workflow for the EcMcrBC complex. (**d**) Local resolution map for the EcMcrBC half-complex. (**e**) Gold-standard FSC curve for the EcMcrBC half-complex after correction for masking effects. The resolution was estimated based on the FSC = 0.143 criterion. (**f**) Cross-validation FSC curves for the EcMcrBC half-complex: cyan curve, refined model *versus* half map 1 used for refinement (Work); red curve, refined model *versus* half map 2 not used for refinement (Free); black curve, refined model *versus* the combined final map (Full). The similarity of the 'work' and 'free' curves suggests no substantial over-fitting. The correlation is above 0.5 up to a resolution of 3.4 Å. (**g**) Cryo-EM densities for selected regions in the map of the EcMcrBC complex.

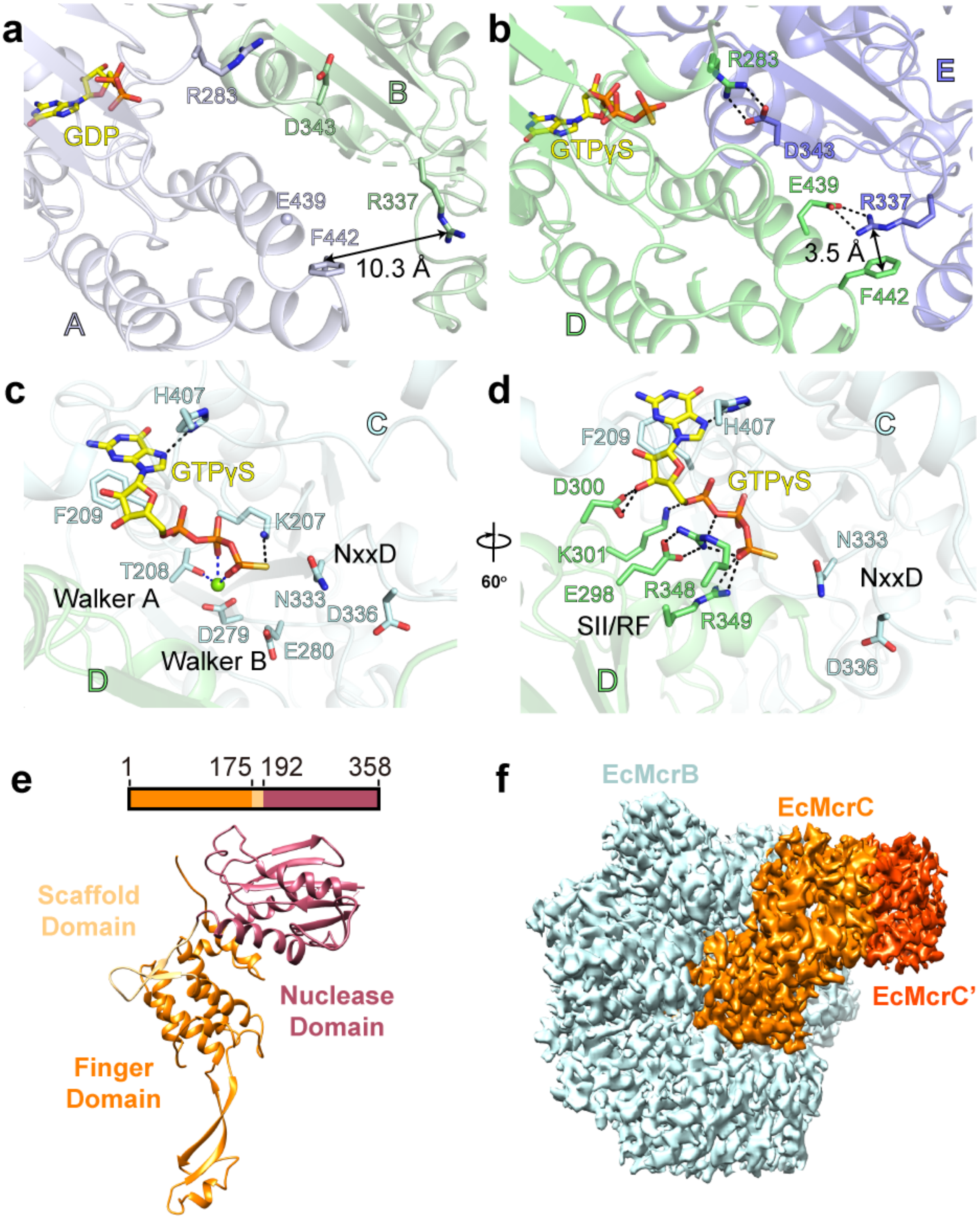


**Figure S5**

**Figure S5: Cryo-EM structure of the EcMcrBC complex, related to Figure 5.** (**a** and **b**) Close-up views of interacting residues at the loose A/B interface (**a**) and the tight D/E interface (**b**) of the

EcMcrB hexamer. Black dashed lines indicate hydrogen bonds. (**c** and **d**) Close-up views of the GTP-binding site at the tight C/D interface, highlighting residues involved in *cis* interactions, in particular those of the Walker A and B motifs and the NxxD motif (**c**), and residues involved in *trans* interactions, in particular those of the Sensor II/arginine finger (SII/RF) motif (**d**). (**e**) Domain architecture of EcMcrC. (**f**) Cryo-EM density map showing the interface between the two EcMcrC subunits (orange and red) and one EcMcrB hexamer (cyan).

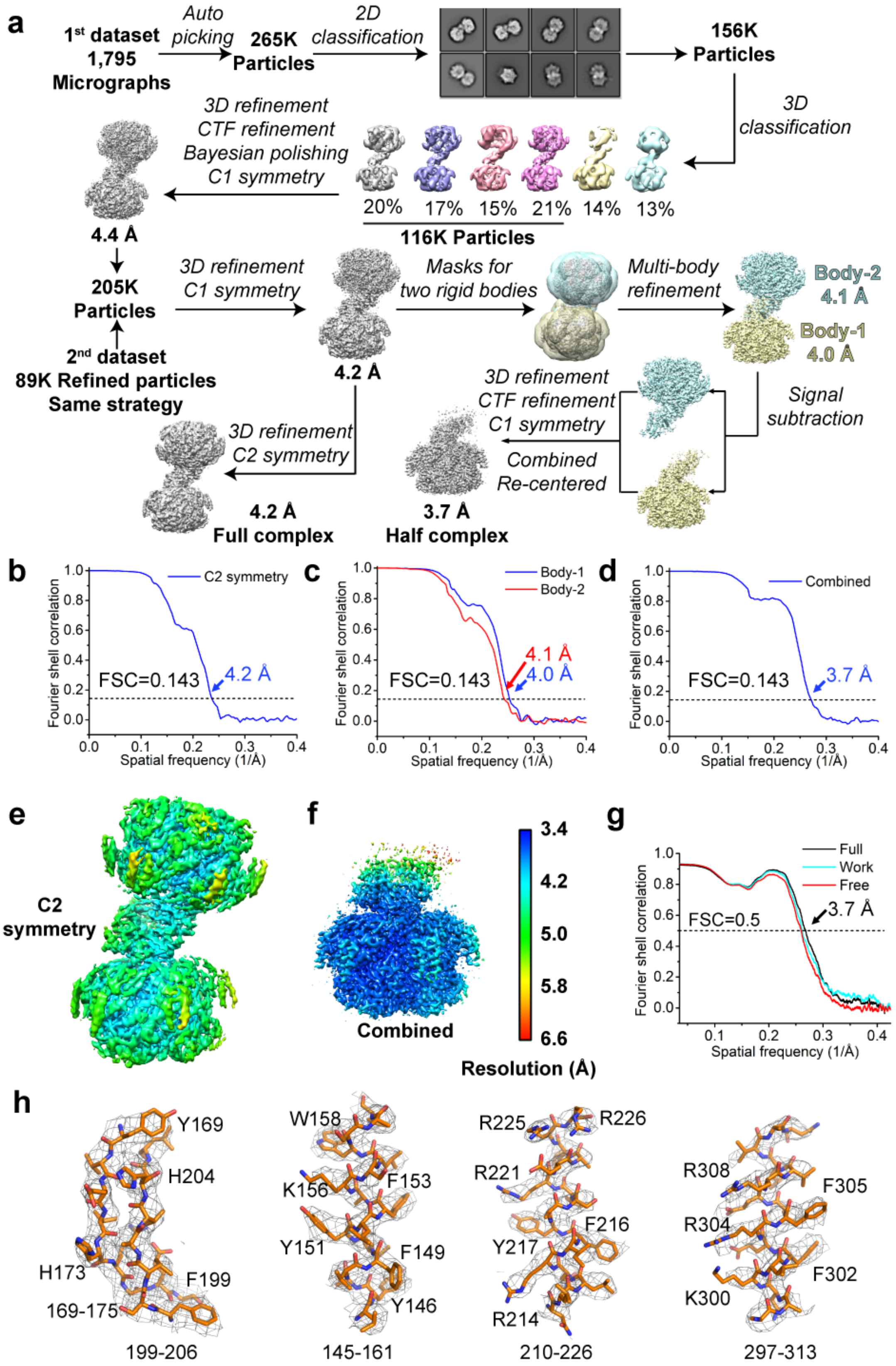
a
1st dataset 1,795 Micrographs
Auto picking
265K Particles
2D classification
156K Particles
3D classification
20% 17% 15% 21% 14% 13%
116K Particles
3D refinement
CTF refinement
Bayesian polishing
C1 symmetry
4.4 Å
205K Particles
2nd dataset
89K Refined particles
Same strategy
3D refinement
C1 symmetry
4.2 Å
Masks for two rigid bodies
Multi-body refinement
Body-2 4.1 Å
Body-1 4.0 Å
Signal subtraction
3D refinement
CTF refinement
C1 symmetry
Combined
Re-centered
3.7 Å
Half complex
3D refinement
C2 symmetry
4.2 Å
Full complex
b
C2 symmetry
FSC=0.143
4.2 Å
Fourier shell correlation
Spatial frequency (1/Å)
c
Body-1
Body-2
4.1 Å
4.0 Å
FSC=0.143
d
Combined
FSC=0.143
3.7 Å
e
C2 symmetry
f
Combined
3.4
4.2
5.0
5.8
6.6
Resolution (Å)
g
Full
Work
Free
FSC=0.5
3.7 Å
h
Y169
H204
H173
F199
169-175
199-206
W158
K156
F153
Y151
F149
Y146
145-161
R225
R226
R221
F216
Y217
R214
210-226
R308
F305
R304
F302
K300
297-313

1 **Figure S6**

**Figure S6: Cryo-EM analysis of the TgMcrB$^{AAA}$C complex, related to Figure 6.** (**a**) Cryo-EM data-processing workflow for the TgMcrB$^{AAA}$C complex. (**b–d**) Gold-standard FSC curves for the maps of the TgMcrB$^{AAA}$C complex obtained with different image-processing strategies after correction for masking effects: (**b**) the map obtained when C2 symmetry was imposed; (**c**) the maps of the two bodies obtained when multi-body refinement was used; and (**d**) the map obtained when the two half-complexes were combined. The resolution was estimated based on the FSC = 0.143 criterion. (**e** and **f**) Local resolution for the cryo-EM map with C2 symmetry imposed (**e**) and the map for the combined half-complexes (**f**). (**g**) Cross-validation FSC curves for the map of the combined TgMcrB$^{AAA}$C half-complexes: cyan curve, refined model *versus* half map 1 used for refinement (Work); red curve, refined model *versus* half map 2 not used for refinement (Free); black curve, refined model *versus* the combined final map (Full). The similarity of the 'work' and 'free' curves suggests no substantial over-fitting. The correlation is above 0.5 up to a resolution of 3.7 Å. (**h**) Cryo-EM densities for selected regions in the map of the combined TgMcrB$^{AAA}$C half-complexes.

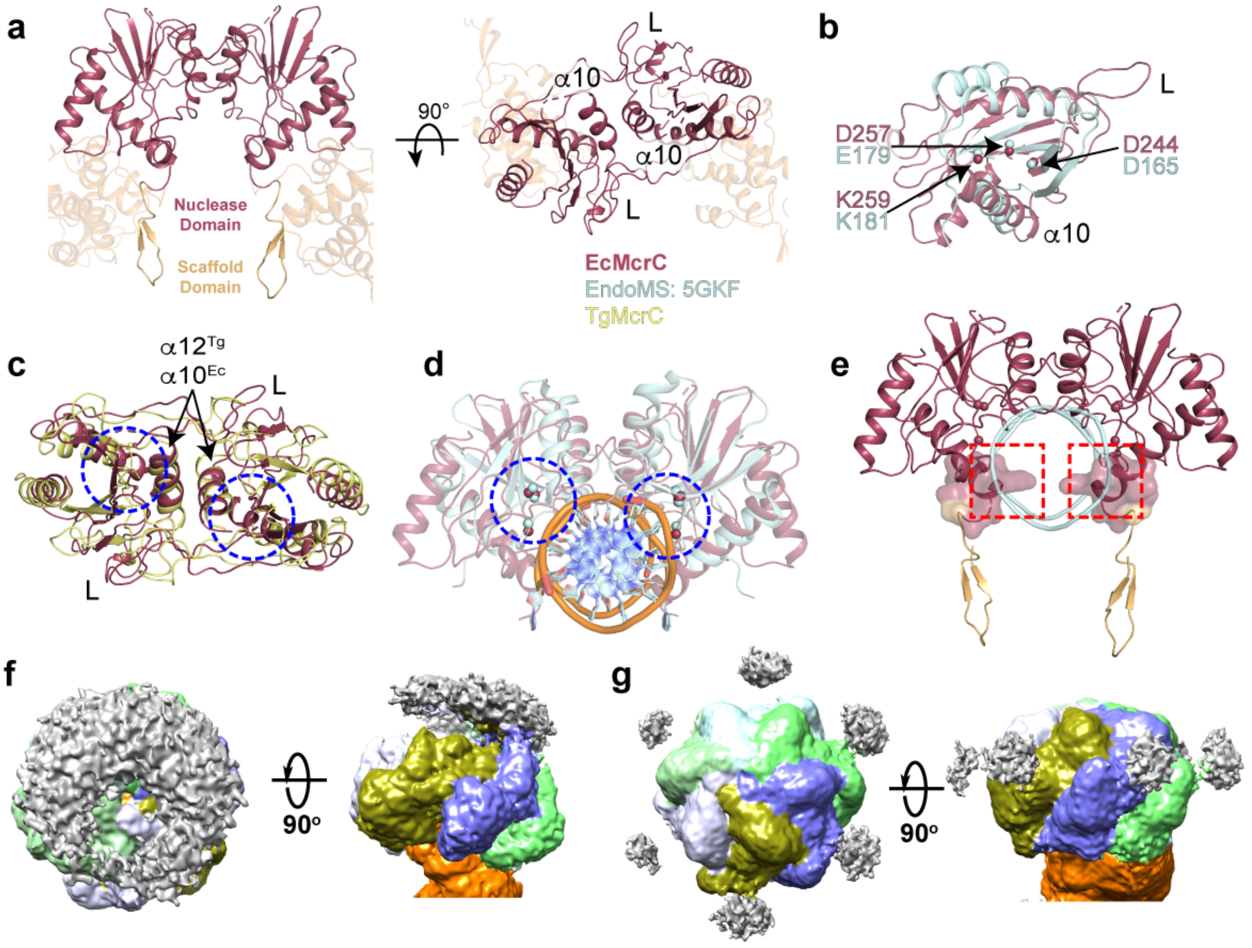


### Figure S7

**Figure S7**: **Structural comparison of the nuclease domains, related to Figure 6.** (**a**) Cartoon representation of the dimer interface of EcMcrC. (**b**) Structural comparison of an EcMcrC monomer with an EndoMS monomer (PDB: 5GKF), showing that the active-site residues overlap (shown as spheres and labeled). (**c**) Superposition of the nuclease domain in the TgMcrC dimer (yellow) and EcMcrC dimer (red). The active sites for DNA cleavage are highlighted by blue circles. (**d**) Structural comparison of the EcMcrC nuclease-domain dimer in the EcMcrBC complex (dark red) with the EndoMS dimer in a DNA-bound state (PDB: 5GKF) (cyan). The active sites for DNA cleavage are highlighted by blue circles. (**e**) Illustration of the cleavage-incompetent conformation of EcMcrC. For clarity, the EndoMS structure is not shown. The backbone of the DNA substrate bound to EndoMS is colored cyan. The red squares indicate regions of potential steric clashes. (**f** and **g**) Low-pass filtered maps of the EcMcrBC complex (**f**) and the TgMcrBC complex (**g**). Unassigned densities are colored in grey and likely represent the six N-terminal DNA-binding domains of the McrB hexamer.

**Supplementary Table S1. X-ray data collection and refinement statistics.**

| | TgMcrB$^{AAA}$ Native | TgMcrB$^{AAA}$ Se derivative |
|---|---|---|
| **PDB ID** | 6UT3 | |
| **Data collection** | | |
| X-ray Source | NECAT 24ID-E | NECAT 24ID-C |
| Wavelength (Å) | 0.9791 | |
| Space Group | $P2_1$ | |
| Cell dimensions | | |
| *a*, *b*, *c* (Å) | 100.02, 108.55, 118.43 | 100.24, 108.87, 118.67 |
| α, β, γ (°) | 90, 106.94, 90 | 90, 107.41, 90 |
| Resolution (Å) | 114.650-2.95 (3.04-2.95) | 113.23-3.14 (3.25-3.14) |
| $R_{\mathrm{merge}}$ (%) | 7.3 (141.5) | 10.6 (138.6) |
| $R_{meas}$ (%) | 7.9 (167.4) | 11.1 (149.5) |
| $CC_{1/2}$ (%) | 99.9 (55.6) | 99.9 (73.8) |
| $I / \sigma I$ | 14.2 (1.0) | 17.3 (1.9) |
| Completeness (%) | 99.6 (99.8) | 99.7 (99.9) |
| Redundancy | 6.8 (6.9) | 13.5 (14.0) |
| **Phasing** | | |
| Initial F.O.M. | | 0.499 |
| Number of sites | | 19 |
| **Refinement** | | |
| Resolution (Å) | 114.65-2.95 | |
| No. reflections | 53,315 (4,037) | |
| $R_{\mathrm{work}}$ / $R_{\mathrm{free}}$ (%) | 34.5/36.4 | |
| No. atoms | | |
| Protein | 15915 | |
| Ligand/ion | 132 | |
| *B*-factors | | |
| Protein | 128.5 | |
| Ligand/ion | 110.4 | |
| Clash score | 13.4 | |
| R.m.s. deviations | | |
| Bond lengths (Å) | 0.007 | |
| Bond angles (°) | 1.21 | |
| Ramachandran plot | | |
| Favored (%) | 94.4 | |
| Allowed (%) | 5.6 | |
| Outliers (%) | 0 | |

*Values in parentheses are for highest-resolution shell. Each dataset was derived from a single crystal.

### Supplementary Table S2. Cryo-EM data collection and refinement statistics.

| | TgMcrB$^{AAA}$ | TgMcrBC | EcMcrBC | TgMcrB$^{AAA}$C (Full mask) | TgMcrB$^{AAA}$C (Body 1) | TgMcrB$^{AAA}$C (Body 2) | TgMcrB$^{AAA}$C (Combined) |
|---|---|---|---|---|---|---|---|
| **EMDB ID** | EMD-20865 | EMD-20866 | EMD-20867 | EMD-20868 | EMD-20869 | EMD-20870 | EMD-20871 |
| **PDB ID** | 6UT4 | 6UT5 | 6UT6 | 6UT7 | | | 6UT8 |
| **Data collection** | | | | | | | |
| Microscope | Titan Krios | | | | | | |
| Detector | K2 summit | | | | | | |
| Voltage (kV) | 300 | | | | | | |
| Pixel size (Å) | 0.50 | | | | | | |
| Total electron exposure ($e^-/Å^2$) | 80.0 | | | | | | |
| Defocus range (μm) | -1.5 to -3.0 | -1.5 to -2.5 | -1.0 to -3.0 | -2.0 to -3.5 | | | |
| Micrographs collected | 1,599 | 2,078 | 1,161 | 4,271 | | | |
| **Reconstruction** | | | | | | | |
| Final particle images | 139,306 | 226,846 | 106,684 | 204,593 | | | 409,186 |
| Pixel size (Å) | 1.25 | 1.0 | 1.0 | 1.25 | | | |
| Box size (pixels) | 256 | 400 | 400 | 256 | | | |
| Resolution (Å) (FSC = 0.143) | 3.14 | 2.44 | 3.28 | 4.26 | 3.95 | 4.10 | 3.68 |
| Map Sharpening B-factor (Å) | -51.1 | -20.6 | -45.2 | -160.4 | -138.2 | -154.8 | -126.9 |
| **Model composition** | | | | | | | |
| Non-hydrogen atoms | 19,126 | 22,374 | 16,938 | 45,062 | | | 22.531 |
| Protein residues | 2,315 | 2,697 | 2,078 | 5,522 | | | 2,761 |
| Water | 12 | 28 | 0 | 0 | | | 0 |
| Ligands | 6 | 6 | 6 | 12 | | | 6 |
| Metals | 6 | 6 | 4 | 12 | | | 6 |
| **Refinement** | | | | | | | |
| Model-to-map CC (mask) | 0.84 | 0.82 | 0.78 | 0.68 | | | 0.80 |
| Model-to-map CC (volume) | 0.82 | 0.81 | 0.77 | 0.72 | | | 0.77 |
| R.m.s deviations | | | | | | | |
| Bond length (Å) | 0.004 | 0.005 | 0.005 | 0.004 | | | 0.004 |
| Bond angles (°) | 0.740 | 0.780 | 0.819 | 0.832 | | | 0.831 |
| **Validation** | | | | | | | |
| MolProbity score | 1.58 | 1.48 | 1.74 | 1.68 | | | 1.66 |
| Clash score | 5.89 | 5.90 | 7.39 | 5.38 | | | 5.08 |
| Ramachandran plot | | | | | | | |
| Outliers (%) | 0.0 | 0.0 | 0.0 | 0.0 | | | 0.0 |
| Allowed (%) | 3.8 | 2.9 | 4.8 | 5.8 | | | 5.8 |
| Favored (%) | 96.2 | 97.1 | 95.2 | 94.2 | | | 94.2 |
| Rotamer outliers (%) | 0.15 | 0.17 | 0.57 | 0.44 | | | 0.44 |
| C-beta deviations (%) | 0.05 | 0.00 | 0.05 | 0.00 | | | 0.00 |
